# An affordable non-invasive validated machine-aided phenotyping pipeline identifies phenotypic variation of stress resilience in alkaline calcareous soil across the life cycle in *Arabidopsis thaliana*

**DOI:** 10.1101/2025.03.02.641020

**Authors:** Marie Knopf, Petra Bauer

**Author notes:** **Author contributions:** Conceptualization (P.B., M.K.), Methodology (M.K.), Investigation (M.K.), Data Analysis (M.K.), Writing - Original Draft (M.K.), Review & Editing (all authors), Supervision (P.B.), Funding Acquisition (P.B.).

## Abstract

Alkaline calcareous soils (ACS) are prevalent globally and challenge plant growth by limiting nutrient uptake, such as iron. The model plant *Arabidopsis thaliana* thrives in disturbed urban environments wherein ACS conditions frequently occur. Existing research largely focused on vegetatively grown *A. thaliana*, while there is a notable lack of studies examining phenotypic variations across the life cycle in ACS. A valuable tool for understanding plant stress resilience is machine-aided phenotyping as it is non-invasive, rapid and accurate. But it is often unavailable to individual plant labs. Here, we established and validated an affordable MicroScan with PlantEye-based machine-aided phenotyping approach, collected and correlated quantitative growth data across plant life cycles in response to ACS.

We used *A. thaliana* wild type and the chlorotic coumarin-deficient mutant *f6’h1-1* to assess weekly morphological and leaf color data both manually and using a multispectral PlantEye device. Through correlation analysis, we selected machine parameters to differentiate size and leaf chlorosis phenotypes. The correlation analysis indicated a close connection between rosette size and multiple spectral parameters, highlighting the importance of the rosette size for plant growth. Most reliable phenotyping was at the beginning bolting stage. This methodology further is validated to detect novel leaf chlorosis phenotypes of known iron deficiency mutants across growth stages.

This affordable machine-aided phenotyping procedure is suitable for high-throughput accurate screening of small-grown rosette plants, such as *A. thaliana*, and enables the discovery of novel genetic and phenotypic variation during the life cycle for understanding plant resilience in challenging soil environments.

**Short summary sentence:** A PlantEye machine-aided non-invasive accurate and reliable phenotyping pipeline depicted the importance of the rosette size for phenotyping and detected leaf chlorosis phenotypes of *A. thaliana* mutants across the life-cycle on alkaline calcareous soil.

**Highlights and major findings:**

- A MicroScan PlantEye machine-aided non-invasive phenotyping pipeline was established for assessing growth data of *A. thaliana* across the life cycle on alkaline calcareous soil and distinguishing leaf chlorosis phenotypes.
- Rosette size was found an important trait that characterizes *A. thaliana* growth.
- Machine phenotyping was most reliable at the beginning bolting stage.
- New phenotypes were detected for Fe homeostasis mutants.

## Introduction

More than 30% of the earth’s soils are alkaline and/or calcareous (Chen & Barak, 1982), and the proportion may rise with global warming and by human activities (Rengel, 2011; Sun et al., 2023). These soils impose several challenges for plant growth (Taalab et al., 2019), and hence it is important to understand which genetic factors help plants to better thrive.

In calcareous soils, calcium carbonate (CaCO_3_) determines the main soil properties. Such soils are formed over calcareous parent rock or by irrigation with carbonate-rich water and often occur in arid or semiarid regions (Wahba et al., 2019). The pH is typically high and buffered by carbonates between 7.5 to 8.2 (Maulood et al., 2012; Wang et al., 2020). Besides calcium (Ca), sodium (Na) is a relevant ion in calcareous soils and additionally involved in alkalinization (Hu et al., 2021). Soils in urban environments are typically alkaline and saline, especially in industrial and traffic areas, but also near residences (Horváth et al., 2015; Yang & Zhang, 2015). *A. thaliana* inhabits many edaphically different habitats, including urban, alkaline and saline environments, indicating it has adaptive genetic mechanisms (Pérez-Martín et al., 2022; Schmitz et al., 2024; Terés et al., 2019). These are not yet fully understood.

Iron (Fe) is in lack in ACS conditions as the bioavailability of Fe is low under high pH (Vélez-Bermúdez & Schmidt, 2023). Iron homeostasis mutants are known in *A. thaliana*, and they often have stronger leaf chlorosis phenotypes than wild type on ACS (Lei et al., 2020; Li et al., 2016; Long et al., 2010; Schmid et al., 2014; Zhang et al., 2015). A key regulator for Fe acquisition is the essential FER-LIKE IRON DEFICIENCY-INDUCED TRANSCRIPTION FACTOR (FIT), and the severely growth-compromized loss of function mutant *fit-3* only grows upon Fe fertilization (Jakoby et al., 2004; Schwarz & Bauer, 2020). One of the FIT target genes encodes FERULOYL-COA 6’-HYDROXYLASE 1 (F6’H1) catalyzing the first enzymatic step in coumarin biosynthesis (Schmid et al., 2014). Coumarins are important secondary compounds that allow *A. thaliana* to mobilize Fe and grow on ACS (Gautam et al., 2021; Long et al., 2010; Robe et al., 2021; Schmid et al., 2014; Terés et al., 2019). *f6’h1* loss of function mutant plants have slight Fe deficiency and leaf chlorosis in alkaline conditions (Robe et al., 2021; Schmid et al., 2014). POPEYE (PYE) is another transcription factor (Long et al., 2010), and the E3 ligases BRUTUS-LIKE1 and 2 (BTSL1/2) interact with transcription factors related with PYE and affect Fe utilization negatively (Lichtblau et al., 2022; Rodríguez-Celma et al., 2019). Loss of function mutants of *pye* and *btsl1 btsl2* do not show leaf chlorosis but complete their life cycles in turf soils without Fe fertilization. Their phenotypes become visible in Fe-limited or resupply conditions (Long et al., 2010; Rodríguez-Celma et al., 2019; Stanton et al., 2023). Many studies on Fe homeostasis in plants focused primarily on the investigation of seedlings on agar plates or in hydroponic solutions containing low Fe concentrations (e.g.(Nguyen et al., 2022; Tabata et al., 2022) or they investigated germination on ACS (Wala et al., 2022).

Plant phenotyping is a powerful method to discriminate genetic diversity across the life cycle, particularly using non-invasive automated methods (Arvidsson et al., 2011; Vasseur et al., 2018). This has been performed in large high-throughput phenotyping pipelines including automatic moving and watering of the plants (Arend et al., 2016; Granier et al., 2006; Pieruschka & Schurr, 2019). These platforms enable non-destructive, fast, standardized, and repeated measurements of single plants over time and can replace tedious manual plant phenotyping and destructive measurements (Poorter et al., 2023). In multispectral scanning, 3D laser scanning provides physical information of an object and positions, while spectral analysis gives hints on plant characteristics like chlorophyll content, nutrient status and water stress (Xia et al., 2023). The laser and near-infrared measurement do not affect photosynthetic performance of the scanned plants (Kjaer & Ottosen, 2015). Despite the advantages of large phenotyping facilities, less automated and smaller lab-based imaging devices would be greatly of benefit to small individual research units. They are affordable solutions, available for restricted budgets and spaces and can be flexibly used. One commercially available, non-destructive phenotyping device is the PlantEye (Vadez et al., 2015) offered in the small and portable format of a MicroScan (Phenospex, Heerlen, The Netherlands). It is an environmental light-independent 3D multispectral imaging system.

Despite of a number of studies on growth of *A. thaliana* in ACS conditions, the phenotypes determined were mainly manually measured phenotypes (Gautam et al., 2021; Long et al., 2010; Schmid et al., 2014; Terés et al., 2019). However, in our view, there is no systematic study addressing the accuracy of machine phenotyping for small rosette plant species like *A. thaliana* in stressful environments causing leaf chlorosis, such as ACS. The PlantEye was used in studies of plant or plant community performance (Gedif et al., 2023; Li et al., 2022; Manavalan et al., 2021; Yang et al., 2022; Zieschank & Junker, 2022). *A. thaliana* was to our knowledge only used in one of them (Yang et al., 2022), but no validation of the approach shown. It is not clear whether machine phenotyping with a PlantEye device discriminates weak leaf chlorosis phenotypes of *A. thaliana* as they occur in ACS.

This study aimed to establish an affordable innovative high-throughput machine-aided phenotyping procedure using the PlantEye suited to detect genetic variation in standard laboratory stress resilience experiments in non-destructive manner throughout the life cycle of *A. thaliana*. Being able to apply machine phenotyping throughout the plant life cycle will be useful for uncovering novel genetic diversity for adaptation in *A. thaliana* to stress conditions like ACS, which could explain the success of this species in urban environments.

## Materials and Methods

### Plant Material

Lines of *Arabidopsis thaliana* (L) Heynh. were multiplied in parallel at Heinrich Heine University. They were wildtype (WT, Col-0) and four mutants in Col-0 background, *f6’h1-1* (Schmid et al., 2014), *fit-3* (Jakoby et al., 2004), *pye-1* (Long et al., 2010) and *btsl1 btsl2* (Rodríguez-Celma et al., 2019).

### Plant growth

Detailed description of plant growth is provided in Supplemental Methods and Supplemental Figure 1. Three plant growth experiments were conducted (Supplemental Figure 1A, B). Briefly, seeds were surface-sterilized, stratified in darkness at 4°C and germinated on ½ upright Hoagland plates for eight days (16 h light, in average 135 µmol·m^-2^·s^-1^, 21 °C in light, 19 °C in darkness and 50% relative humidity). On the eighth day, plants were transferred to soil. Pots filled with soil were covered with matt blue vinyl foil, leaving a hole for plants to grow (Supplemental Figure 2A). Plants were grown in plant cabinets with 16 h light (98-112 µmol·m^-2·^s^-1^, Polyklima True Daylight + LED), 21 °C during light, and 19 °C during darkness. Humidity was not controlled. Eight pots fitted in one tray. Soil was prepared with a peat substrate supplemented with different amounts of CaCO_3_, NaHCO_3_, and sand according to Supplemental Figure 2B and as specified in the text, named control, ACS1-5, and ACS3-25% and -50% sand. The pH was determined according to the below procedure. In experiment 3, plants were moved to a walk-in growth chamber with 16 h light (80-120 µmol·m^-2·^s^-1^, BX120c4, VAYOLA, Helsinki, Finland), 21°C day temperature, 19 °C night temperature and 57% humidity after three weeks in soil instead of being kept in a growth cabinet. 16 plants were grown per line and condition for experiments 1 and 2, eight for experiment 3 (Supplemental Figure 1A, B). Trays were regularly rotated.

### Soil pH determination

Deionized water was added to 15 g wet soil (5g soil dry weight) to reach 50 ml in a falcon tube, rotated for 30 min with 20 rotations per min and centrifuged for 10 min at 4000g and 20 °C (Heraeus Multifuge X1R, Thermo Fischer Scientific, Hampton, USA). The supernatant was filtered through paper filter (Folded filters, 322345, Schleicher&Schuell, Dassel, Germany) and the pH was determined with a pH electrode (S20 Seven Easy, Mettler Toledo, Columbus USA). An average was calculated from two samples per condition.

### Destructive determination of the chlorophyll content

The chlorophyll content was determined from whole rosettes of plants. The rosettes were frozen and grinded in liquid nitrogen, and ca. 100 mg plant material was used, weighed and an acetone extraction procedure used. After centrifugation at 15,000*g* for 10 min, the absorption was measured at 470 nm, 642 nm and 662 nm (Shimadzu UV visible Spectrophotometer UVmini-1240, Duisburg, Germany and Hellma OS 104-OS cuvette, Müllheim, Germany) and pigment contents per fresh weight calculated according to the following formulas (Lichtenthaler, 1987). Chlorophyll a: (µg/ml) = (11.24*A662 - 2.04*A642) * dilution; Chlorophyll b: (µg/ml) = (20.13*A642 - 4.19*A662) * dilution; Carotenoids: (µg/ml) = [(1000*A470-1.90*Chla - 63.14*Chlb)/214] * dilution.

### Manual morphological phenotyping

Plants were photographed. Different parameters were manually determined either using the photos and performing measurements with Image J or by directly measuring plants. Manual measurements were performed to determine per plant the rosette diameter [cm], manual rosette area [mm^2^], manual rosette convex hull [mm^2^], plant fresh weight [mg], rosette fresh weight [mg], plant height [cm], number of siliques, number of side branches and flowering time [days after sowing, DAS] as outlined (Supplemental Figure 3; details in Supplemental Methods).

### Machine (PlantEye) phenotyping

Machine phenotyping was performed with a MicrosSan device with PlantEye F600 (Phenospex, Heerlen, The Netherlands) and the implemented Software HortControl (Phena version 2.0, HortControl version 3.85). Automated phenotyping requires improved techniques to exclude non-plant background from analysis (Arvidsson et al., 2011; Li et al., 2014; Vasseur et al., 2018). Plant pots were therefore covered with blue foil and measured one by one by placing the pots in a blue holder that kept the plants at a fixed height and within the measured unit. Each plant was either measured four times (experiments 1 and 2), two times (Experiment 3) or once (experiment 2, pigment content extraction), at each time point for technical replication of the measurement. Plant pots were turned by 90° between the repeated measurements. The color range from Hue values between 200-360° (blue to purple) was removed from all images that were used for calculations to remove non-plant background. The PlantEye parameters were split into morphological parameters (Supplemental Figure 4) and spectral parameters (Supplemental Figure 5). The morphological parameters were the 3D Leaf Area [mm^2^] (three dimensional area of the plants), Canopy Light Penetration Depth [mm] (how far laser reaches into the canopy), Convex Hull Area [mm^2^] (Area of a convex hull drawn around the plant), Convex Hull Area Coverage [%] (proportion of the Convex Hull overlaid by the plant), Convex Hull Maximum Width [mm] (longest straight length in the Convex Hull), Convex Hull Aspect Ratio [%] (Quotient of the Convex Hull Maximum Width and the perpendicular line to it at its midpoint), Convex Hull Circumference [mm], Plant Height Max [mm] (Distance from the pot height to the highest point of the plant), Plant Height Averaged [mm] (highest point of the plant is replaced by the average height of the highest 10% of points in the image), Projected Leaf Area [mm^2^] (area covered by plant seen from above), Digital Biomass [mm^3^] (product of 3D Leaf Area and Plant Height Averaged), Surface Angle Average [°] (average angle of all triangles between points forming a plant) and Voxel Volume Total [mm^3^] (sum of all voxels representing the plant). Using non-invasive phenotyping plant leaf color shades can be distinguished (Dobbels & Lorenz, 2019; Matsuda et al., 2012; Ochogavía et al., 2014). The spectral parameters were Hue Average [°] (color value of the HSL, Hue-Saturation-Lightness color space), Saturation Average [%] (saturation value of HSL, grey 0% to pure color 100%), Lightness average [%] (lightness value of HSL, white 100% to black 0%), Greenness Leaf Index (GLI, (2·GREEN-RED-BLUE)/(2·GREEN+RED+BLUE)), Normalized Difference Vegetation Index (NDVI, (NIR-RED)/(NIR+RED)), Normalized Pigment Chlorophyll Index (NPCI, (RED-BLUE)/(RED+BLUE)) and Plant Senescence Reflectance Index (PSRI, (RED-BLUE)/NIR). For all these parameters the average per plant and the percentage of voxels within different (definable) ranges were calculated in the HortControl software implemented in the PlantEye using the pre-set values.

### Statistical analysis

For the comparison of several groups, two-way ANOVA and Tukey Test were performed in R (R 4.4.0). Figures were prepared in Statistical Package for Social Science (SPSS, International Business Machines Corporation, version 29.0.0.0, licence version 5725-A54) or Microsoft Office 2016 Excel. Correlation analysis was done in SPSS. After test for normal distribution (Kolmogorov-Smirnov) Spearman-Rho correlation and graph plotting were done in SPSS. Hierarchical clustering was done in R (R 4.4.0). The data were split between time points to avoid disturbance of the analysis by non-available data. Data were z-score transformed by phenotypic parameters (scaler function, liver package, R). Then graphs were created using the dist (method: euclidean) and hclust (agglomeration method: complete) function (stats package).

## Results

### Correlation Analysis of Manually and Machine-Derived Phenotypic Parameters in Plant Growth on Alkaline-Calcareous Soil

Automated phenotyping requires improved techniques to exclude non-plant background from analysis (Arvidsson et al., 2011; Li et al., 2014; Vasseur et al., 2018), while plant leaf color shades can be distinguished (Dobbels & Lorenz, 2019; Matsuda et al., 2012; Ochogavía et al., 2014). The strain of natural calcareous soil conditions can be imitated (Ben Abdallah et al., 2017; Busoms et al., 2023; Ding et al., 2020; Gautam et al., 2021; Msilini et al., 2009; Murgia et al., 2015; Pérez-Martín et al., 2022; Rosenkranz et al., 2021; Schmid et al., 2014; Terés et al., 2019). Much of the published phenotypic screening in *A. thaliana* has been conducted at the seedling or early vegetative stage (DeLoose et al., 2024; Satbhai et al., 2017; Schmid et al., 2014), precluding observations of phenotypic diversity during reproduction. Hence, not all Fe deficiency phenotypes have been fully exploited. Machine-driven phenotyping would clearly be a benefit. The MicroScan system with the PlantEye device (Phenospex, Heerlen, The Netherlands) is affordable and hence is a small, portable and cost-effective solution for plant phenotyping in a single lab setting. While not fully automated, the system allows rapid and frequent plant scanning, enabling time course data collection over the life cycle of individual plants with minimal effort.

Here, we set out to validate this system for its application potential for small rosette plant species and its effectiveness in high-throughput phenotyping under alkaline calcareous soil (ACS) conditions. We assumed that specific manually measured phenotypic parameters previously recorded in ACS could be replaced by PlantEye parameters, offering the many advantages associated with machine phenotyping, such as increased objectivity, efficiency, and reduced labor intensity (Akhtar et al., 2024). At first, we collected suitable manual and machine-derived plant growth data for *A. thaliana* wildtype (WT) and its coumarin-deficient mutant *f6’h1-1* under control and up to seven ACS conditions, representing a scale of differing pH values from pH 6.2 (control) up to in between 7.8 (intermediate ACS) and 8.3 (severe ACS) (details in Supplemental Figure 2B). Data recording was performed throughout the growth cycle in two independent experiment series (overview in Supplemental Figure 1A; all plant growth phenotyping data in Supplemental Table 1, details on parameters in Supplemental Figures 3-5). Importantly, the *f6’h1-1* mutant indeed suffered more severe iron deficiency and chlorosis than the wildtype under alkaline conditions, as expected (Schmid et al., 2014). In a further preliminary experiment, we had confirmed that indeed growth in an ACS condition causes reduced size of WT and *f6’h1-1* with more intense visible leaf chlorosis in *f6’h1-1*, confirmed by manual measurements of rosette diameter and SPAD values (Supplemental Figure 6).

Then, we used these growth data and conducted a correlation analysis between 12 manually measured and 20 PlantEye-derived parameters. We detected many correlations in the datasets (Figure 1), which was a strong hint on the accuracy of the phenotyping procedure. Very importantly, each of the manual parameters matched with at least four or more significantly correlating PlantEye values. Key findings included correlation of manual parameters like rosette diameters, plant height, plant weight and chlorophyll content with their PlantEye equivalents like 3D Leaf Area, Plant Height Max, Digital Biomass and spectral parameters including Hue Average, PSRI Average, NDVI Average and Lightness Average. Surprisingly, one manual parameter, namely the rosette diameter, significantly correlated with all 20 PlantEye parameters. All manual parameters related to rosette size (in addition to rosette diameter also manual rosette area, manual rosette convex hull and rosette weight) correlated best with 20, 19, 19 and 17 PlantEye parameters. This impressive finding speaks in favor of the importance of the rosette size for *A. thaliana* growth physiology and development.

**Figure 1:**
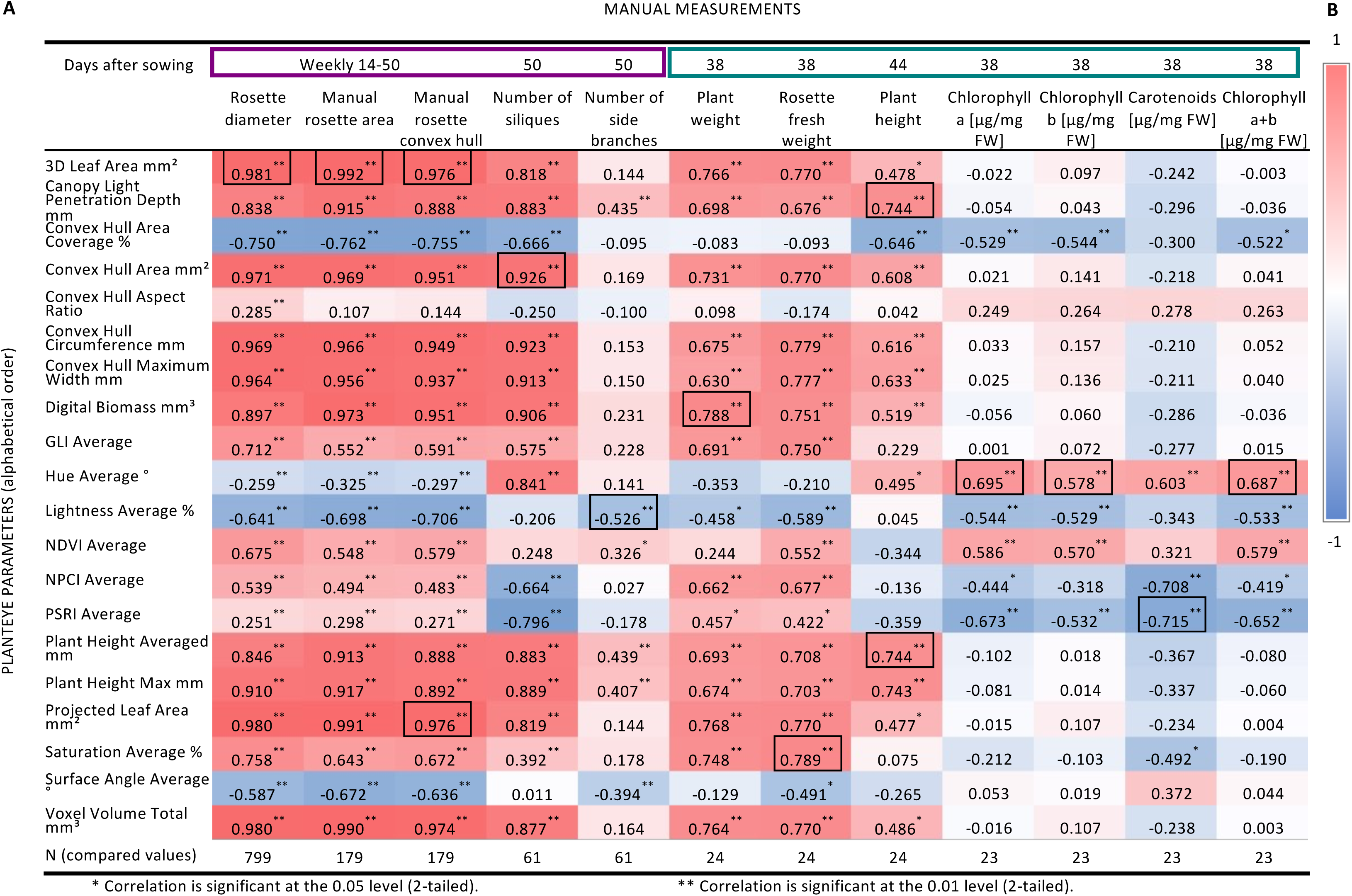
Correlation Analysis of Manually and Machine-Derived Phenotypic Parameters. **A:** Data of twelve manual and 20 machine-derived PlantEye parameters for A*. thaliana* wildtype (WT) and its coumarin-deficient mutant *f6’h1-1* under control and up to seven alkaline calcareous soil (ACS) conditions, representing a scale of differing pH values from pH 6.2 (control) up to 8.3 (severe ACS), recorded during two experiments and subjected to correlation analysis. Data were collected at the indicated time points. Rectangles around time points indicate two experiments (lavender: experiment 1; purple: experiment 2). As indicated, either rosettes (pigment content, rosette fresh weight) or whole plants (remaining parameters) were used. Spearman Rho analysis was conducted in SPSS (* p<0.05, ** p <0.01). Heatmap color codes correlation coefficients (-1 blue, 0 white, +1 red). The strongest correlation per manual parameter is marked with a box. N= number of data points as indicated. **B:** Color scale of correlation coefficients (-1 blue, 0 white, +1 red).

To study accuracy in detail, we investigated three rosette size correlations using scatterplot analysis of the data points (Figure 2A-C). Initially during the growth cycle, the strong correlation between rosette diameter and 3D Leaf Area (correlation coefficient 0.981) was rather square (R^2^ = 0.934) than linear (R^2^=0.887). However then, scattering of the data points became stronger starting 28 days after sowing (Figure 2A), when flowering began (Supplemental Figure 7). Second, the manual rosette area correlated very strongly with the machine 3D Leaf Area (correlation coefficient 0.992). This correlation was linear (R^2^ =0.964) up to 42 days after sowing (Figure 2B). Third, for the manual rosette convex hull, the strongest correlation was not found with the machine Convex Hull Area (correlation coefficient 0.951), but with the machine 3D Leaf Area and the machine Projected Leaf Area (correlation coefficient 0.976) (Figure 1). This can again be explained by an effect of inflorescence stem formation on the measurements especially beyond 41 days after sowing, as indicated by the increased scattering at those time points (Figure 2C). Hence, the correlations are best at the rosette growth stage up to the bolting stage but diminish at the advanced reproductive stages with emerged and branched inflorescence stems and siliques. ACS treatments did not affect flowering time in the two experiments (Supplemental Figure 7A, B).

**Figure 2:**
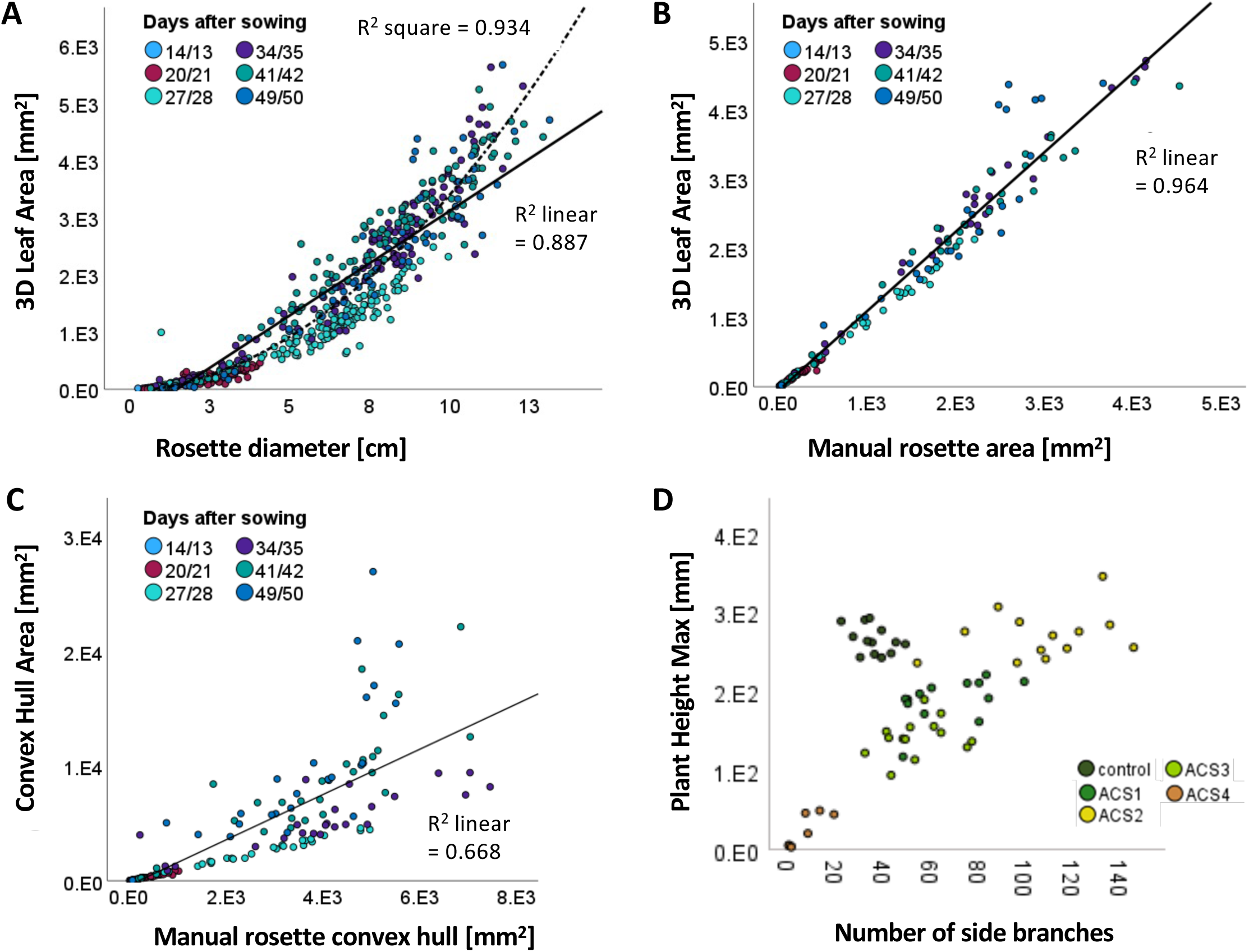
Scatterplots Analysis of Selected Manually and Machine-Derived Phenotypic Parameters. **A, B, C:** Scatter plots of rosette diameter with PlantEye 3D Leaf Area (N=799), manual rosette area with PlantEye 3D Leaf Area (N=179) and manual rosette convex hull with PlantEye Convex Hull Area (N=179). Growth data for *A. thaliana* wildtype (WT) and its coumarin-deficient mutant *f6’h1-1* under control and alkaline calcareous soil (ACS) conditions, representing a scale of differing pH values from pH 6.2 (control) up to 8.3 (severe ACS) recorded at six time points, color coded in the scatter plots (14/13, 20/21, 27/28, 34/35, 41/42 and 49/50 days after sowing). **D**: Manual number of side branches with PlantEye Plant Height Max (N= 61), 50 days after sowing. Colors indicate different growth conditions, control and varying ACS conditions from ACS1 to ACS4 (details in Supplemental Figure 2). Scatter plots were created in SPSS.

Observations during flowering led us to the next interesting question, whether flowering time would be correlating with any PlantEye parameters and also whether it is among the traits, that can be estimated with the PlantEye. We subjected manual flowering time data to correlation analysis with all PlantEye data collected throughout the life cycle at various time points. Notably, the three strongest (negative correlations) were found with Canopy Light Penetration Depth, Plant Hight Averaged and Plant Height Max 28 days after sowing (Supplemental Figure 7C, D) coinciding with the average flowering time of 27 days after sowing (Supplemental Figure 7), revealing that for the estimation of flowering time, PlantEye measurements close to flowering time are needed. As flowering time and plant height are meaningful parameters reflecting reproductive growth progression during the life cycle of *A. thaliana,* we also analyzed the plant height. Plant Height Averaged and the Light Canopy Penetration Depth (correlation coefficient 0.744) (Figure 1) correlated equally well with the manual plant height, making both suitable to replace manual height measurements.

Next, we addressed the question whether the Digital Biomass was indeed the best parameter representing the plant weight in *A. thaliana*. We analyzed plants with and without inflorescences stems, imitating two growth stages. Surprisingly, while the whole plant weight indeed correlated best with the Digital Biomass (correlation coefficient 0.788), the rosette fresh weight correlated better with seven other PlantEye parameters (Figure 1). The Digital Biomass was therefore the most suitable estimator for *A. thaliana* weight only after inflorescence stems were formed, while other PlantEye parameters were even more suitable for non-flowering plants.

Surprisingly, manual parameters related to reproductive success (number of siliques and side branches) had different correlations with PlantEye parameters. While the number of siliques correlated with 16 out of 20 PlantEye parameters, it was only six for the number of side branches (Figure 1). Those included Lightness Average, Plant Height Averaged, Canopy Light Penetration Depth, Plant Height Max, Surface Angle Average and the Normalized Difference Vegetation Index (NDVI) in that order of correlation strength. The correlation with the height parameters can be due to longer plants having more side branches, but the negative correlation with the Lightness remains unexplained. The comparably weak correlation of the number of side branches with the PlantEye parameters makes it less suitable for being replaced. A possible explanation could be that not all side branches were detected in the measurement, or the number side branches simply is not in a relation with overall plant growth and physiology in the tested growth conditions. To further look into that, we had a look at the data of the Plant Height Max and the side branches (Figure 2D). This revealed a better correlation if only the ACS plants and not the control plants were considered indicating an effect of the condition on the correlation strength. Strikingly, only a single PlantEye parameter correlated by far least with any manual parameters, which was Convex Hull Aspect Ratio (Figure 1), indicating that this parameter is not suitable for the described purpose.

Due to the particular importance of chlorosis as symptom in ACS, we were especially interested, which PlantEye parameters would be useful for estimating chlorophyll content in *A. thaliana.* This was determined in the rosettes of 38-day-old plants after their inflorescence stems were removed. The correlations of rosette leaf plant pigmentation were consistent but surprisingly not very high (Figure 1). The strongest correlation for Chlorophyll a+b content was found for with the Hue, PSRI, NDVI and Lightness (correlation coefficients 0.687, -0.652, 0.579 and -0.533). There were not only correlations with the averages of spectral parameters, but also their bin values, further confirming their usefulness for chlorophyll estimation (Supplemental Table 2).

Finally, we were very intrigued by the many correlations among various parameters. Some closely related parameters reflected similar traits. We figured that it should be possible to investigate relatedness of parameters by hierarchical clustering of manual and machine phenotyping parameters. Indeed, this approach highlighted clustering of corresponding parameters (Figure 3). Overall, manual parameters clustered closely together. Yet, the manual parameters often clustered best with corresponding machine parameters. For example, the manual rosette size parameters (rosette diameter, manual rosette area, manual rosette convex hull) were best clustering with all corresponding morphological machine parameters such as Plant Height Max, Convex Hull Area, Convex Hull Circumference, Convex Hull Maximum Width, Voxel Volume Total, 3D Leaf Area and Projected Leaf Area (Figure 3A). Similar situations were seen for plant weight and plant height (Figure 3B, C). Remarkably, the chlorophyll a, b and a+b contents clustered closest with the NDVI Average and the carotenoid content with the Hue Average (Figure 3D). On the other hand, as suggested by the results of the correlation analysis, the number of siliques and number of side branches were not clustering closely together, and only distantly clustering machine parameters were identified (Figure 3E).

**Figure 3:**
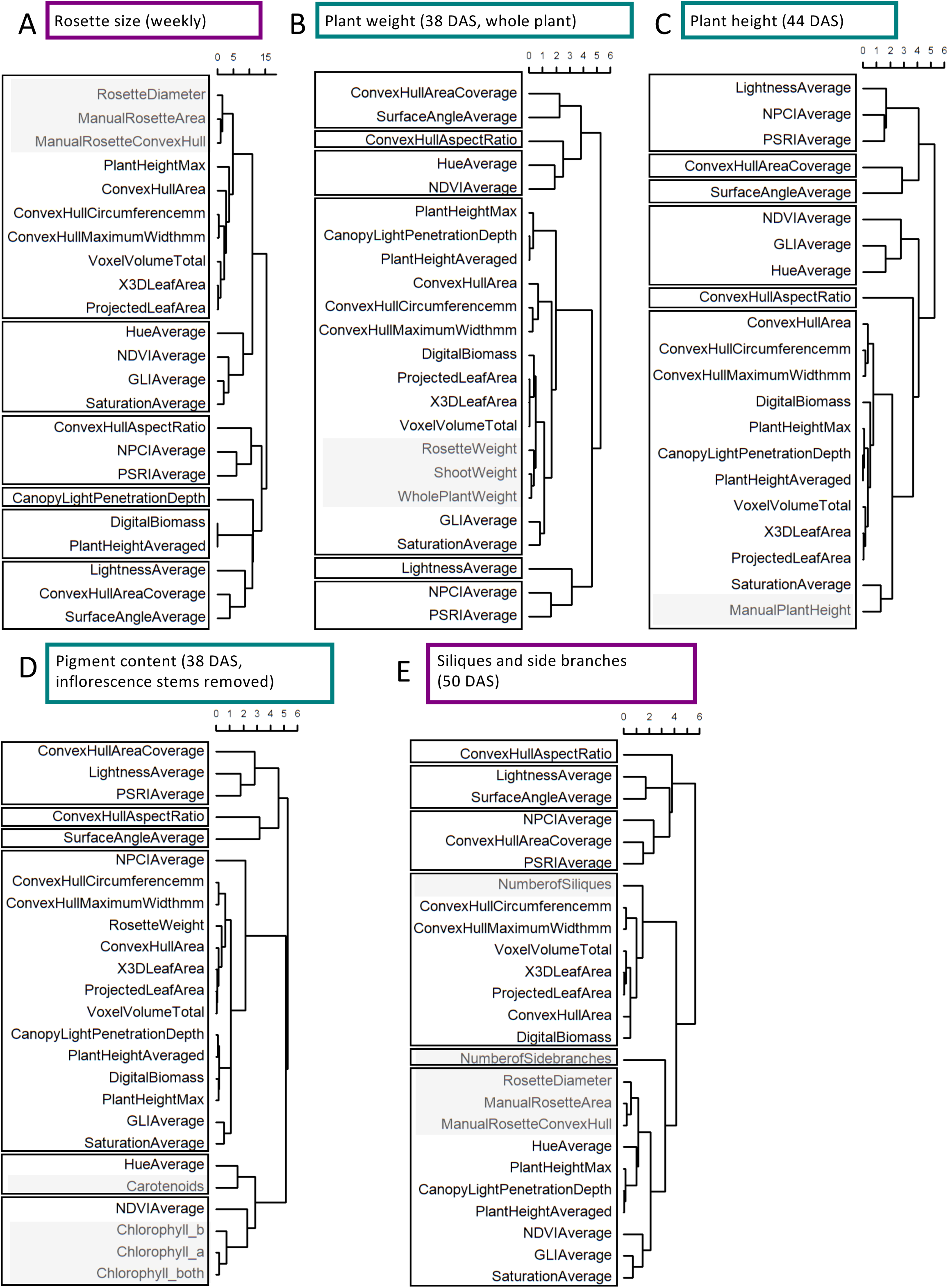
Hierarchical Clustering of Manually and Machine-Derived Phenotypic Parameters. *A. thaliana* wildtype (WT) and the coumarin deficient mutant *f6’h1-1 were* grown in seven alkaline calcareous soil (ACS) conditions with varying pH values in two experiments to collect manual and machine-derived (PlantEye) data. Manual parameters were clustered with machine parameters collected at the same time points. **A**: Weekly determined rosette size parameters (rosette diameter, manual rosette area, manual rosette convex hull), **B:** plant weight 38 days after sowing (DAS) (rosette weight, shoot weight and whole plant weight), **C:** manual plant height 44 DAS **D:** rosette pigment content 38 DAS (chlorophyll a, chlorophyll b, chlorophyll a+b, carotenoids) and **E:** number of siliques and side branches 50 DAS. Hierarchical clustering was done using the hclust function in R (stats package). Colored rectangles around boxes indicate two different experiments (experiment 1: purple, experiment 2: green). Manually determined parameters are marked with a grey box. Splitting into six clusters is highlighted by black rectangles.

In conclusion, machine-aided phenotyping with PlantEye was found to deliver accurate and reliable data that were meaningful for growth physiology in *A. thaliana* plants grown in ACS conditions. We identified optimal machine-aided parameters, such as 3D Leaf Area, Hue Average, NDVI Average, PSRI Average and Lightness Average covering well the manually phenotyped traits. A best time point for reliable machine phenotyping of *A. thaliana* was upon around the onset of inflorescence stem elongation at days 34-35. Rosette size was found to be a meaningful plant trait, correlating with morphological and plant color traits. Number of siliques was more suited for correlation with machine parameters than the number of side branches.

### Assessment of Machine-aided Phenotypic Analysis of Wild Type and the coumarin-deficient Mutant *f6’h1-1* in Seven Artificially Created ACS Conditions

Having established and validated machine phenotyping, we then inspected closely the obtained plant data at days 27-28 and 34-35, focusing on the best suited machine-aided phenotyping parameters to assess differences in growth physiology between WT and *f6’h1-1* plants in the ACS conditions described above and in experiments 1 and 2 (Supplemental Figure 1A, 2). Our aim was to identify one ACS condition, that distinguishes best WT and *f6’h11* plants with machine phenotyping and that still allows plants to complete their life cycles. We predicted that this was the case for an intermediate ACS condition. Having such an ACS condition combined with meaningful PlantEye parameters will allow us in the future investigating genetic adaptation to ACS in this species in clearly defined soil conditions with reduced time expense for phenotyping.

When we displayed the plant phenotyping data (Figure 4; Supplemental Figure 8), we excluded strong ACS conditions because plants had died at 28 days after sowing (DAS) (Figure 4A; Supplemental Figure 8A), and mild ACS due to the lack of phenotypes (Figure 4C – E, Supplemental Figure 8C - G) except for the number of siliques in *f6’h1-1* (Figure 4F). ACS-induced small growth was detectable at both time points in both lines in ACS3 (*f6’h1-1* at 28 DAS in experiment 2, Figure 4G), ACS4, ACS3-25% sand and ACS3-50% sand (Figure 4C, G, Supplemental Figure 8C, H). Phenotypes were most obvious in ACS4 and ACS3-50% sand. Small growth was more severe in *f6’h1-1* than in WT in ACS3 at 35 days in experiment 2 (Figure 4G). Among the intermediate ACS conditions, leaf color changes were found in ACS3, ACS4, ACS3-25% sand and ACS3-50% sand with differences between *f6’h1-1* and WT (Figure 4 D-E and H-I, Supplemental Figure 8, D-G and I-L). Based on the strong influence on rosette size and its large interquartile range in ACS3-50% sand, we also excluded ACS4 and ACS3-50% sand (Figure 4C and G). Since ACS3-grown *f6’h1-1* plants despite a slightly larger rosette size frequently showed stronger affected spectral parameters than in ACS3-25% sand, we finally opted for ACS3 as a best condition for detecting genetic variation in future experiments (Figure 4H, I, Supplemental Figure 8J).

**Figure 4:**
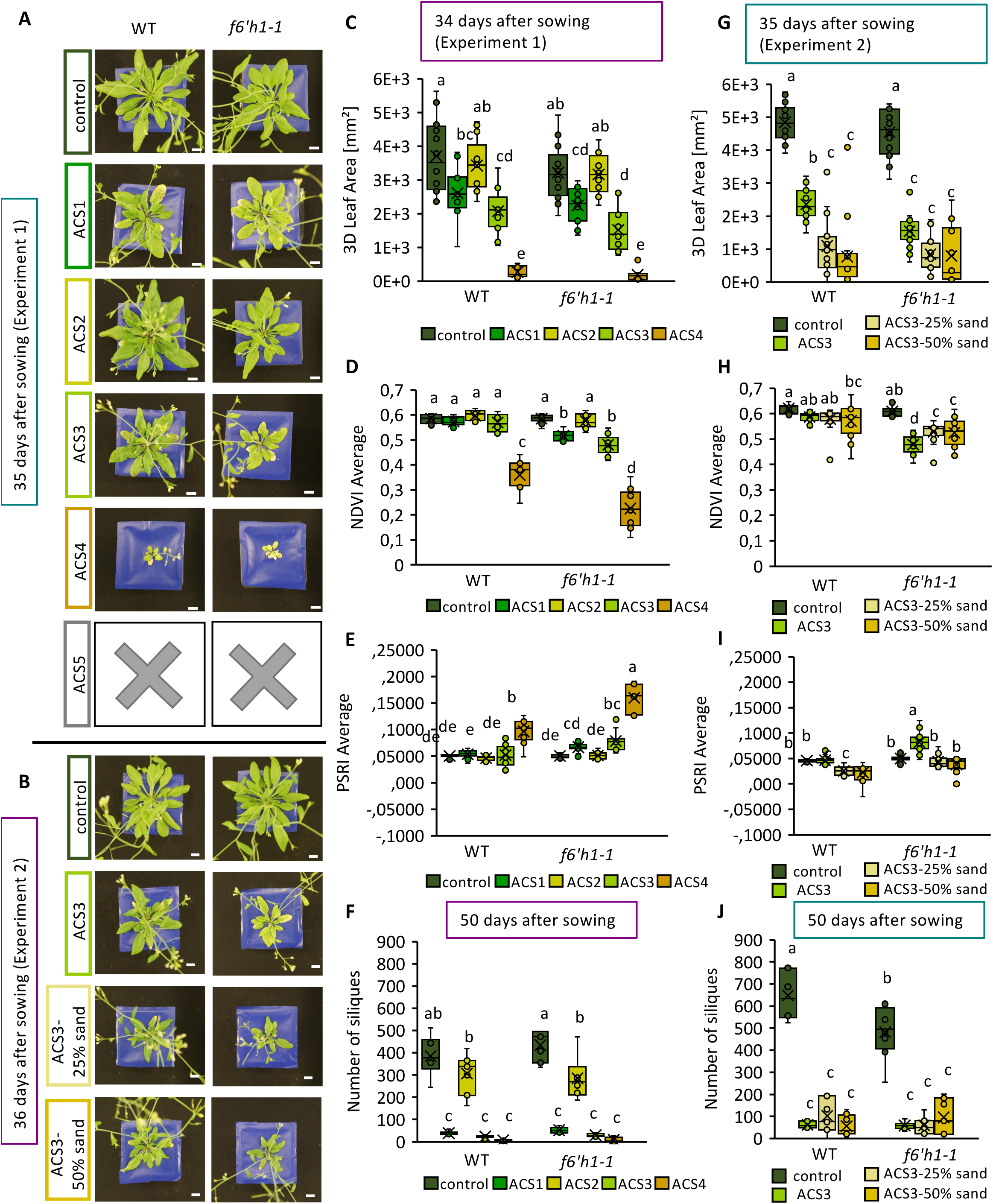
Machine-aided Phenotypic Analysis of Wild Type and the Coumarin-deficient Mutant *f6’h1-1* in Seven Artificially Created ACS Conditions. *A. thaliana* wildtype (WT) and the coumarin deficient mutant *f6’h1-1 were* grown in seven alkaline calcareous soil (ACS) conditions with varying pH values in two experiments to determine an intermediate condition differentiating the phenotypes of lines. Selected phenotypic parameters are shown. **A, B**: Photos of plants in ACS conditions. Scale bar = 1 cm. Plants in ACS5 had died at that time point. **C, D, E, F:** 3D Leaf Area, Normalized Difference Vegetation Index (NDVI) Average, Plant Senescence Reflectance Index (PSRI) Average and number of siliques (manual) of the plants in experiment 1 at 34 and 50 DAS. **G, H, I, J:** 3D Leaf Area, NDVI Average, PSRI Average and number of siliques (manual) of the plants in experiment 2 at 35 DAS and 50 DAS. Letters indicate statistical differences (Two-way ANOVA, Tukey Test in R, p=0.05, N= 9-16 plants).

In sum, the established manual and machine phenotyping system was suited to detect morphological and leaf color changes between control / ACS and WT / *f6’h1-1*, with the intermediate ACS3 condition selected as suitable for detecting genetic variation. This approach provides a clearly defined pipeline for plant growth and non-invasive phenotyping. It facilitates the detection of variation in *A. thaliana* lines with varying performance in ACS or any treatment leading to similar effects. The findings also confirm the importance of coumarins produced with the help of F6’H1 for adaptation to ACS conditions.

### Validation of Manual and Machine-aided Phenotypic Analysis using Known Regulatory Iron Homeostasis Mutants

In the final step, the chosen ACS3 condition and the selected phenotyping parameters were applied to validate the procedure using iron (Fe) homeostasis mutants with the additional aim of identifying potentially novel phenotypes appearing during the life cycle (all phenotyping data in Supplemental Table 3). The mutants were selected to have differing Fe deficiency-induced leaf chlorosis symptoms, namely the severely chlorotic loss of function mutant *fit-3* (Jakoby et al., 2004), the *pye-1* mutant turning chlorotic on ACS (Long et al., 2010), and the Fe-accumulating mutant *btsl1 btsl2* (Hindt et al., 2017; Lichtblau et al., 2022; Rodríguez-Celma et al., 2019). The growth phases and the influences of nutrient deficiencies on the growth cycle have not yet been completely shown for any of the three mutants in quantitative phenotyping experiments in an ACS condition. We expected that the mutants might differ in their growth curves on ACS with regard to wild type revealing novel phenotypes allowing to resolve their functional context during the life cycle.

First, we analyzed the *fit-3* mutant. It showed visibly reduced growth in control and ACS3 condition 14 days after sowing as compared to WT. Reduced NDVI was visible in plant images (Figure 5A) and quantitatively confirmed (Figure 5B). Interestingly, in the *fit-3* mutant, there was no increase in the 3D Leaf Area at all during the experiment in neither condition, while there was a strong increase for the WT from 15 to 42 days after sowing in control and less pronounced in ACS3 (Figure 5C). The halting of rosette growth of *fit-3* clearly shows growth reduction caused by the inability to take up nutrients like Fe in any growth condition. Next, *pye-1* showed a growth curve similar to WT in control condition. Growth in ACS was much reduced compared to the WT (Figure 5D, E) and it also had a reduced NDVI Average at 28 days compared with other lines and conditions (Figure 5F). Strikingly, the *btsl1 btsl2* double mutant remained slightly smaller or similarly large like WT in control and ACS3 condition at earlier growth phases, before it increased its 3D Leaf Area in both conditions between 28 and 35 days after sowing and outgrew the WT (Figure 5D). This was the case at an earlier time point in ACS3 than in control. Therefore, 29 days after sowing, the *btsl1btsl2* plants were smaller than WT in control condition but not in ACS3 (Figure 5D, E). The NDVI was unchanged (Figure 5F).

**Figure 5:**
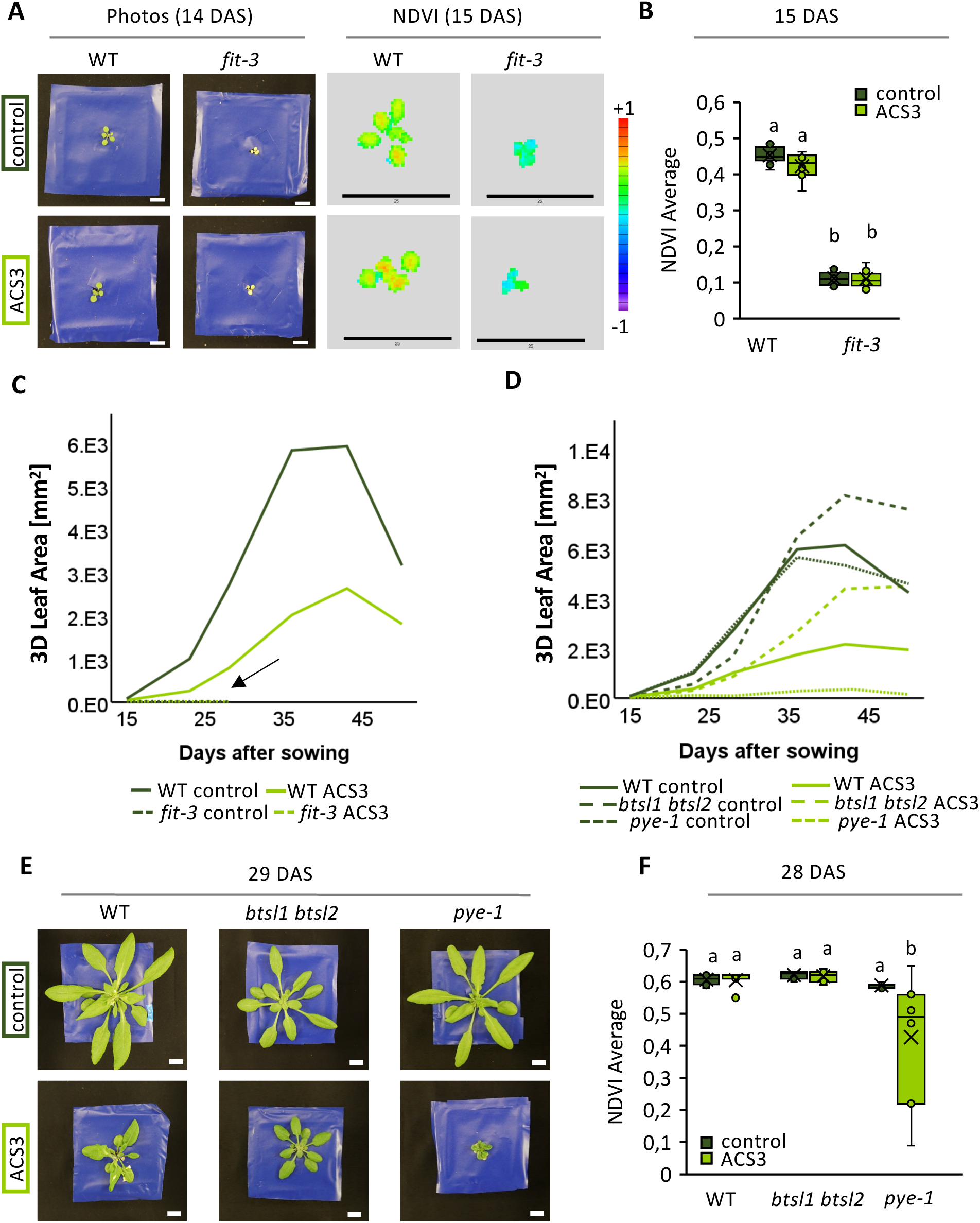
Machine-aided Phenotypic Analysis Using Known Regulatory Iron Homeostasis Mutants. The chosen alkaline calcareous soil (ACS3) condition and the selected phenotyping parameters were applied to validate the procedure using the iron (Fe) homeostasis mutants *fit-3, pye-1* and *btsl1btsl2*. **A:** Photos and Normalized Difference Vegetation (NDVI) images (PlantEye) of wildtype (WT) and *fit-3* mutant 14/15 days after sowing (DAS). Scale bar photos = 1cm, cale bar NDVI images = 25 mm. Scale of the NDVI ranging from purple -1 to red +1. **B:** Quantification of the NDVI. **C:** Development of the 3D Leaf Area of WT and *fit-3* (arrow) in control soil and ACS3 soil over the experiment. N= 4-8 plants. **D:** Development of the 3D Leaf Area of WT, *btsl1btsl2* and *pye-1* over the time of the experiment. N=4-8 plants. **E:** Photos of WT, *btsl1btsl2* and *pye-1* in control and ACS3 at 29 DAS. **F:** NDVI of WT, *btsl1btsl2* and *pye-1* at 28 DAS, N=7-8 plants. Letters indicate statistical differences (Two-way ANOVA, Tukey Test in R, p=0.05, N= 7-8 plants).

In conclusion, Fe-related growth phenotypes of the investigated Fe homeostasis mutants like Fe deficiency-induced growth reduction and leaf chlorosis or in opposite to that possible Fe-related growth increase could be detected with machine phenotyping during the life cycle. This validates the use of rapid, precise and reliable machine phenotyping on ACS3 for investigating genetic adaptation.

## Discussion

This study established reliable and accurate procedures for growing *A. thaliana* in alkaline calcareous soil (ACS) and phenotyping plants using both manual and machine-aided methods across their life cycle. We identified an optimal ACS condition (ACS3), enabling clear differentiation of leaf color changes and growth patterns between WT and Fe homeostasis mutants using machine-aided phenotyping (PlantEye). Clearly, our here-established machine phenotyping pipeline is applicable to support small plant research labs in their endeavors to uncover the numerous molecular-physiological and developmental patterns affecting growth rates in the model reference species *A. thaliana*.

### Machine-aided Phenotyping Across the Life Cycle of *A. thaliana* Plants Growing on ACS is Reliable and Accurate

Reliable phenotyping requires broad phenotype testing, treatment differentiation, and ground-truth validation (Manavalan et al., 2021; Nguyen et al., 2016; Ziamtsov & Navlakha, 2019). PlantEye data effectively distinguished ACS and control plants, matching manual measurements in plant size and leaf color (Figure 4 and Supplemental Figure 6). Notably, the PlantEye performed consistently across two independent experiments (Figure 4D, H), underlining reliability. 3D Leaf Area showed high correlation with ground-truth manual rosette measurements, especially before inflorescence formation (Figure 2A, B). Remarkably, although chlorophyll-spectral parameter correlations were not as strong as for rosette size, pre-bolting and round-bolting measurements (27/28 and 34/35 DAS) were already sufficient to differentiate growth conditions and genotypes based on spectral parameters (Figure 4, Supplemental Figure 8).

### Different Machine Parameters (PlantEye) were Correlating with Manually Determined Parameters

Key PlantEye parameters, such as 3D Leaf Area, accurately replaced manual measurements of rosette diameter and area until 42 days after sowing (Figure 2A, B). Spectral parameters—Hue, NDVI, PSRI and Lightness—correlated with chlorophyll content, reducing the effort for manual assessment. Additional correlations were observed with siliques, plant weight, and flowering time (Figure 1). Rosette diameter strongly correlated with 3D Leaf Area (R² = 0.934), which also correlated with manual rosette area (R² = 0.964) (Figure 2A, B), consistent with similar findings in crops like soybean and peanut (Manavalan et al., 2021; Vadez et al., 2015). To account for diurnal leaf movement (Poorter et al., 2023), we prefer 3D Leaf Area over Projected Leaf Area. Whole plant weight also correlated with Digital Biomass (R² = 0.881) after inflorescence stem formation, similar to wheat and rye studies (Bazhenov et al., 2023).

Notably, while no previous studies linked PlantEye parameters to number of siliques and side branches, we found a correlation between number of siliques and plant area (e.g Convex Hull Area and 3D Leaf Area) that also correlated with the rosette diameter, suggesting a link between rosette size and reproductive fitness (Clauss & Aarssen, 1994). This may be genotype- and environment-dependent and requires further characterization. Side branch correlations were weak, though data hinted at condition-dependent correlations (Figure 2D). Our study showed more side branches in higher control plants than smaller ACS plants. This observation was contradicted a study in melon plants reporting reduced shoot branching under alkaline conditions (Ulas et al., 2019). Further investigation would be needed

Since chlorosis is a key iron deficiency symptom in ACS (Abadía et al., 2011) spectral parameters such as Hue, PSRI, NDVI and Lightness were found as important traits for chlorophyll estimates, though correlations were weaker than those of rosette area measurements. That might be due to their original design not being for single plant chlorophyll estimation (Hassan & Gutub, 2022; Huang et al., 2021; Merzlyak et al., 1999). However, the existence of correlation of Hue, PSRI, NDVI and Lightness with chlorophyll content aligns with previous reports (Castro & Sanchez-Azofeifa, 2008; Chen et al., 2021; Merzlyak et al., 1999; Sass et al., 2012; Tuncay, 2011; Wu et al., 2008), while less correlating indices, like NPCI and GLI, were designed for other questions (Louhaichi et al., 2001; Peñuelas et al., 1994).

Flowering time correlated negatively (R = –0.83 to –0.84) with Canopy Light Penetration Depth and Plant Height just around bolting (Supplemental Figure 7), because plants taller at that time point had flowered earlier. With repeated measurements around the flowering time, the PlantEye could be used to estimate flowering time as inflorescence stem length. Other definition methods of flowering time, like first visible bud or flower opening were not tested but seem more difficult due to higher resolution needed. The Convex Hull Aspect Ratio was unsuitable for *A. thaliana* analysis in ACS indicating that there is no influence on it, but can be involved in other applications like species or growth stage determination (Choudhury et al., 2016; Haque & Haque, 2018).

Ultimately, NDVI, Lightness, Hue, and PSRI proved effective for chlorosis detection, while 3D Leaf Area was the best alternative for manual rosette size estimation, supporting the PlantEye as a valuable tool for machine aided *A. thaliana* phenotyping in ACS.

There were also limitations. While PlantEye data correlated with reproductive traits, the system cannot directly spot siliques or side branches in 3D models, limiting its reliability for silique counts. Moreover, alkaline conditions reduce silique size (Jain & Schmidt, 2024), complicating reproductive fitness assessments. Inflorescence stem growth also introduced measurement inaccuracies, though straightening shoots could improve precision. Nevertheless, 3D Leaf Area measurements remained reliable until 41/42 DAS (Figure 2B, C). The best measurement window was at 34/35 DAS, around a week after bolting, when ACS and control differences were most pronounced (Figure 4; Figure 5C; Supplemental Figure 8), before data started scattering (Figure 2) and inflorescence stems overshadowed the rosette. Future studies should examine correlations between spectral parameters and chlorophyll content over time for improved assessments.

### The Machine Phenotyping of WT and Fe Homeostasis Mutants Detected Expected and New Leaf Chlorosis Phenotypes on a Suited ACS condition

The selected condition ACS3 produced consistent size reduction of WT, one Fe deficiency and one Fe homeostasis mutant, *f6’h1t* and *pye*, comparable to reduced size of *A. thaliana lines* in natural ACS (Terés et al., 2019), and across three experiments (Figure 4 A, B; Figure 5, E). It was shown that upon iron deficiency treatment, rosette growth can arrest (Truong et al., 2024), and especially in young leaves (Ngigi et al., 2024). However, we did not observe a complete arrest in rosette growth of the WT, but rather a reduced growth as observed in natural ACS (Terés et al., 2019). Complete growth arrest was only observable in *fit-3* (Figure 5C) and an tendency in *pye-1* (Figure 5D). This was especially pronounced, between 28 and 35 days, when growth in the control condition was strongest. Future studies could investigate the reasons behind reduced rosette growth and especially the role of iron availability in that. The affected spectral parameters in *f6’h1-1* in ACS aligned with previous reports on chlorosis in alkaline soils (Schmid et al., 2014) similar to some natural lines in natural ACS (Terés et al., 2019). The reproducibility and effect on plant growth make ACS3 a reliable tool for growing and phenotyping *A. thaliana* lines.

The *fit-3* mutant is severely iron deficient (Jakoby et al., 2004) and failed to grow in both control and ACS3 conditions, emphasizing FIT’s crucial role in iron homeostasis overriding any condition effect (Connorton et al., 2017; Schwarz & Bauer, 2020). FIT is a master regulator controlling multiple downstream genes including *F6’H1* and many others performing various reactions in Fe acquisition and metal homeostasis upon Fe uptake in the root (Schwarz et al., 2020). The *pye-1* mutant, previously reported to arrest at the cotyledon stage in alkaline soil (Long et al., 2010), remained growth-impaired even when transplanted to ACS3 at day eight, suggesting persistent sensitivity beyond early development. The leaf chlorosis, seemingly more prevalent in the oldest leaves (Figure 5E, F), may indicate its inability to mobilize Fe or control proper levels of other heavy metals like zinc and manganese. This observation coincides with a previous report on differing roles of old and young leaves in metal ion homeostasis in *A. thaliana* (Ngigi et al., 2024, bioRxiv). The *btsl1btsl2* mutant (Hindt et al., 2017) had not been tested in alkaline calcareous soil before. It initially showed reduced growth but later surpassed WT in both conditions, with an earlier growth recovery in ACS3 (Figure 5D). This is highly interesting as it might suggest a potential advantage of *btsl1btsl2* loss of function in alkaline conditions while maintaining normal growth in iron-sufficient environments. That aligns with previous reports of improved growth in iron-limited conditions without iron toxicity symptoms under iron sufficient conditions (Hindt et al., 2017). In agreement with a functional model proposed by (Lichtblau et al., 2022), the better growth of the *btsl1btsl2* mutant can be explained by reduced downregulation of transcription factors (group IVc including bHLH104, ILR3, bHLH115 and bHLH34) that promote the Fe uptake response when BTSL1 and BTSL2 proteins are not there (Hindt et al., 2017; Lichtblau et al., 2022) (rs). Overexpression of the transcription factors has likely a similar effect as *btsl1btsl2* knockout to stimulate Fe uptake (Liang et al., 2017; Wang et al., 2017; Zhang et al., 2015). Normally, the BTS(L)-type E3 ligase proteins may bind the transcription factors and prepare them for degradation through protein ubiquitination (Rodríguez-Celma et al., 2019; Selote et al., 2014; Spielmann et al., 2023). Clearly, the here-presented phenotyping pipeline is suitable to detect reliably plant phenotypes of novel Fe homeostasis mutants. The study highlights the benefits of multi-stage phenotyping to capture dynamic growth patterns. Future studies should replicate findings for *pye-1* and *btsl1btsl2* mutant interactions and explore additional iron-regulatory mutants to better understand ACS tolerance mechanisms.

A limitation is the possible variability in peat-based soil composition. Using reference lines across experiments can help mitigate this. Uncontrolled humidity and blue foil may have influenced plant responses to water availability. While artificial ACS conditions offer higher controllability and consistency across labs, they do not fully reflect natural alkaline calcareous soil structure. Future studies could explore the comparability of artificial ACS with different natural ACS conditions to further validate the system.

In conclusion, this study outlines the preparation of several ACS conditions, with ACS3 emerging as an effective tool for detecting phenotypic differences in wildtype (Col-0) and Fe homeostasis mutants. Importantly, this growth condition will serve in the future for assessment of growth phenotypes of mutants that help to further depict the intricacies of Fe regulatory mechanisms.

## Conclusions

In conclusion, this study achieved multiple key objectives: (1) demonstrating an affordable, reliable and accurate non-invasive high-throughput phenotyping pipeline to be applied by a single lab working with a small rosette plant species like *A. thaliana*, (2) identifying optimal phenotyping parameters for *A. thaliana* in ACS using PlantEye, (3) defining ACS3 as a condition that effectively differentiates wildtype from *f6’h1-1*, and (4) identifying new growth phenotypes of Fe homeostasis mutants using this established pipeline.

These findings lay a foundation for further research on plant adaptation to alkaline calcareous soils and numerous other stress factors. The successful application of the PlantEye highlights its potential for large-scale non-invasive studies on plant stress responses to ACS opening up new opportunities for future research on *A. thaliana* mutants and natural variation accessions. Beyond that, provided some adjustments, this procedure is easily applicable to any plant species of a size that can be assessed by the MicroScan device with PlantEye.

## Acknowledgements

This work is funded by the Deutsche Forschungsgemeinschaft (German Research Foundation) grant – Project ID 456082119 – TRR 341/1 project A08 to PB. The authors thank Hans-Jörg Mai for advice on statistics and Elke Wieneke for her support in enabling large-scale plant growth. The help of Maren Huppertz and Florian Pollmeyer with plant growth and phenotyping is greatly appreciated. The authors are thankful to members of TRR 341 for suggestions and help with statistics. We also thank the Phenospex Support Team for addressing any technical issues.

During the preparation of this work the authors used ChatGPT 4.0 in order to refine the text for clarity, coherence, and readability in the manuscript preparation process. After using this tool/service, the authors reviewed and edited the content as needed and take full responsibility for the content of the publication.

## Glossary

ACS: Alkaline Calcareous Soil
FIT: FER-LIKE FE DEFICIENCY-INDUCED TRANSCRIPTION FACTOR
F6’H1: FERULOYL-CoA 6′-HYDROXYLASE1
PYE: POPEYE
BTSL: BRUTUS-LIKE
NDVI: Normalized Difference Vegetation Index
PSRI: Plant Senescence Reflectance Index
GLI: Greenness Leaf Index
NPCI: Normalized Pigment Chlorophyll Index

## Supplemental data

**Supplemental Figure 1:**
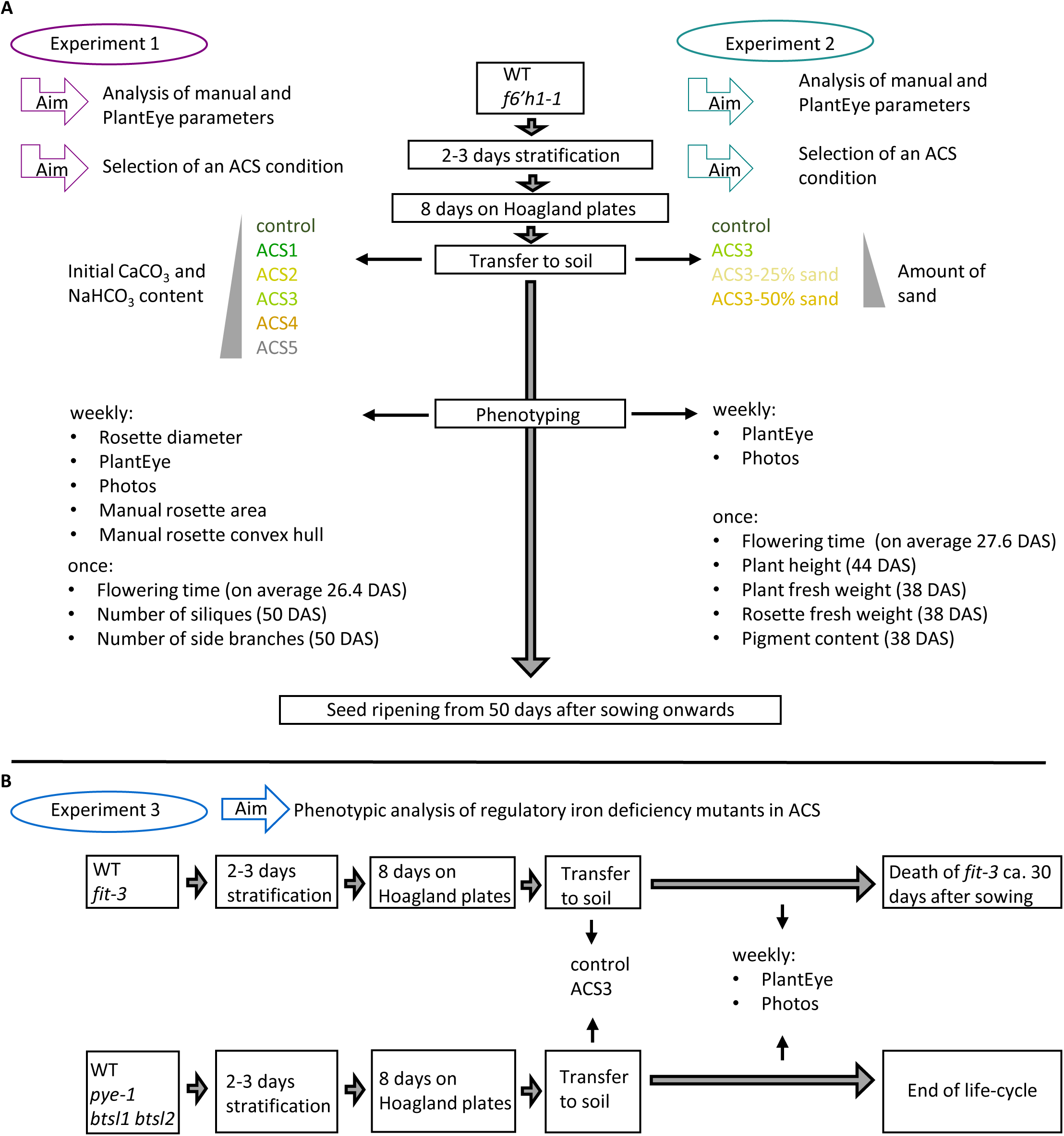
Overview Over Three Experiments for Phenotypic Data Collection and Mutant Analysis. **A:** The wild type (WT) and coumarin deficient mutant *f6’h1-1* were grown in two experiments (Experiment 1 and 2) in mild to severe alkaline calcareous soil (ACS) conditions to collect manual and machine-derived (PlantEye) phenotypic parameters and determine an intermediate alkaline calcareous soil (ACS) condition showing differences between the WT and *f6’h1-1*. ACS conditions were created by adding CaCO_3_, NaHCO_3_ and sand to a peat-based soil substrate resulting in a pH range between 6.2 and 8.3. Phenotypic measurements were conducted throughout the plants’ life cycle. **B:** In experiment 3, wildtype (WT) and three regulatory iron homeostasis mutants (*fit-3, pye-1, btsl1btsl2*) were grown in control and ACS3 and machine-derived parameters were determined to detect potential novel phenotypes.

**Supplemental Figure 2:**
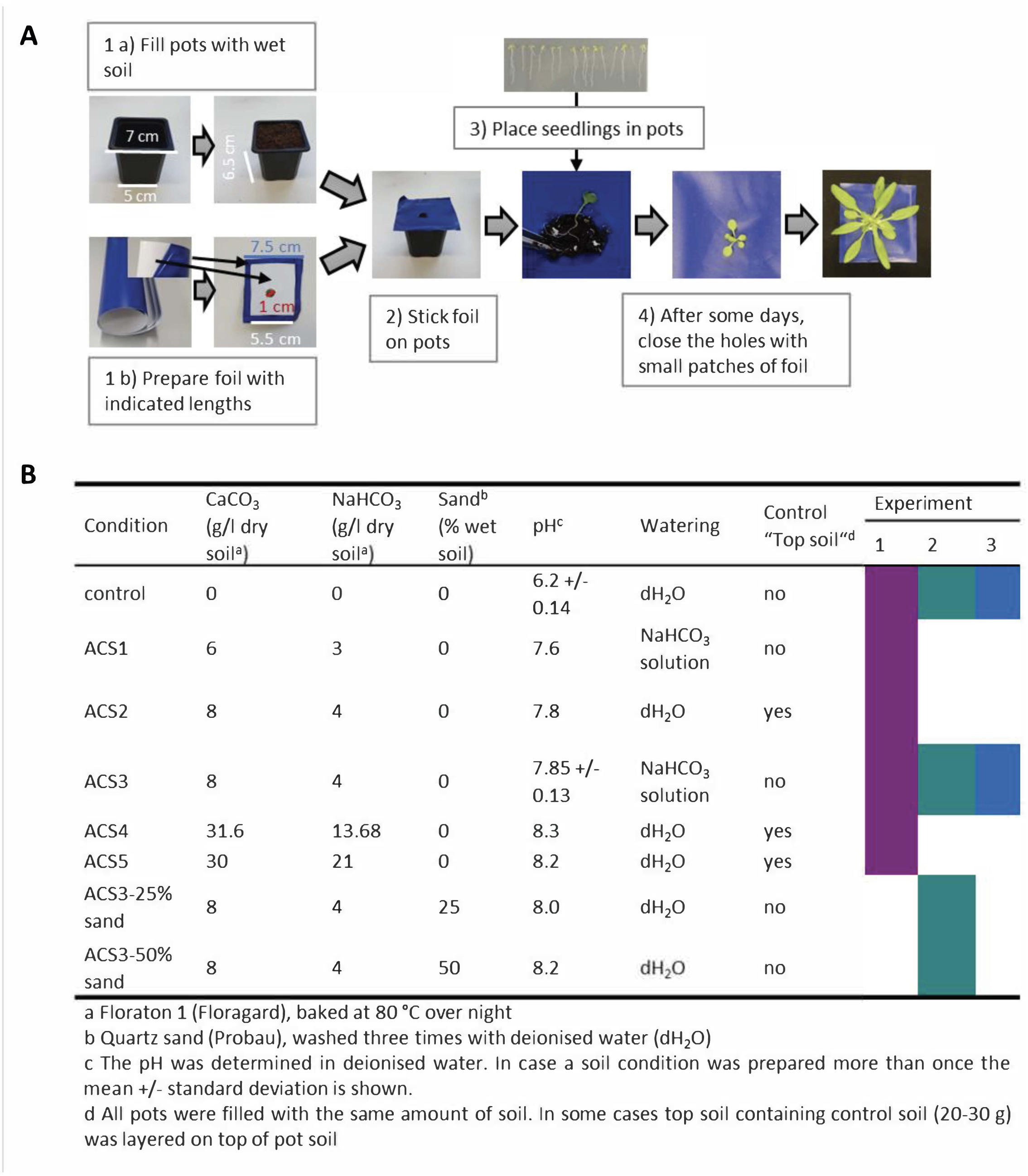
Method of Planting and Preparation of Alkaline Calcareous Soil (ACS). **A:** To facilitate background removal for the machine-aided measurements, soil was covered with a blue foil, before planting, leaving a hole into which eight-day-old seedlings were planted. Holes were then closed around the seedling. **B:** Detailed composition of control and different alkaline calcareous soil (ACS) conditions. Soil was prepared with a peat substrate supplemented with different amounts of CaCO_3_, NaHCO_3_, and sand, named control, ACS1-5, and ACS3-25% and ACS3-50% sand. The pH of the ACS conditions ranged from mild (pH 7.6) to severe (pH 8.3).

**Supplemental Figure 3:**
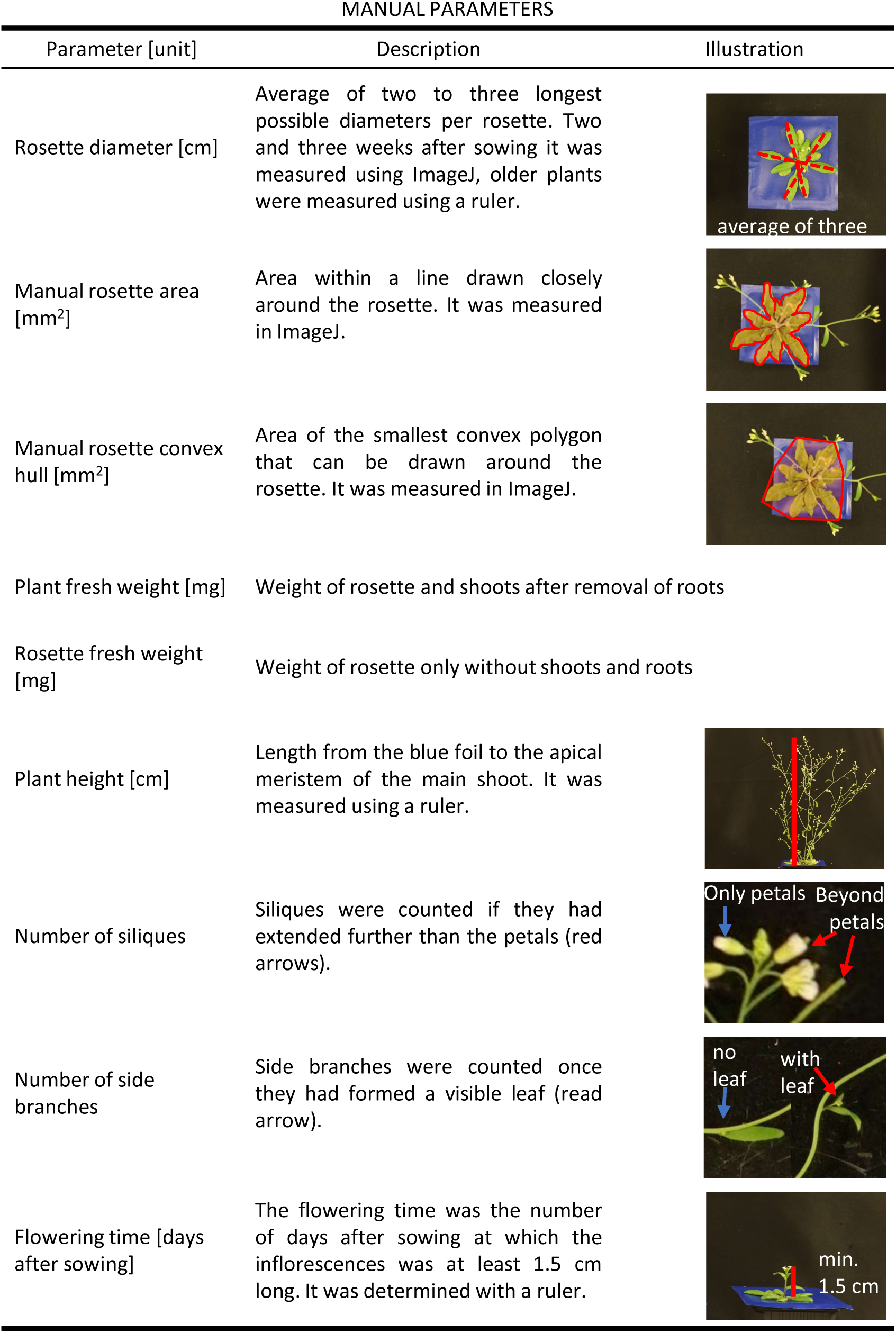
Manually Measured Phenotypical Parameters. The phenotypical parameters were either determined using photos and ImageJ or directly at the plants using a ruler. The rosette diameter, rosette area and rosette convex hull were determined weekly, the plant weight 38 days after sowing (DAS), the plant height at 44 DAS and the number of siliques and side branches at 50 DAS.

**Supplemental Figure 4:**
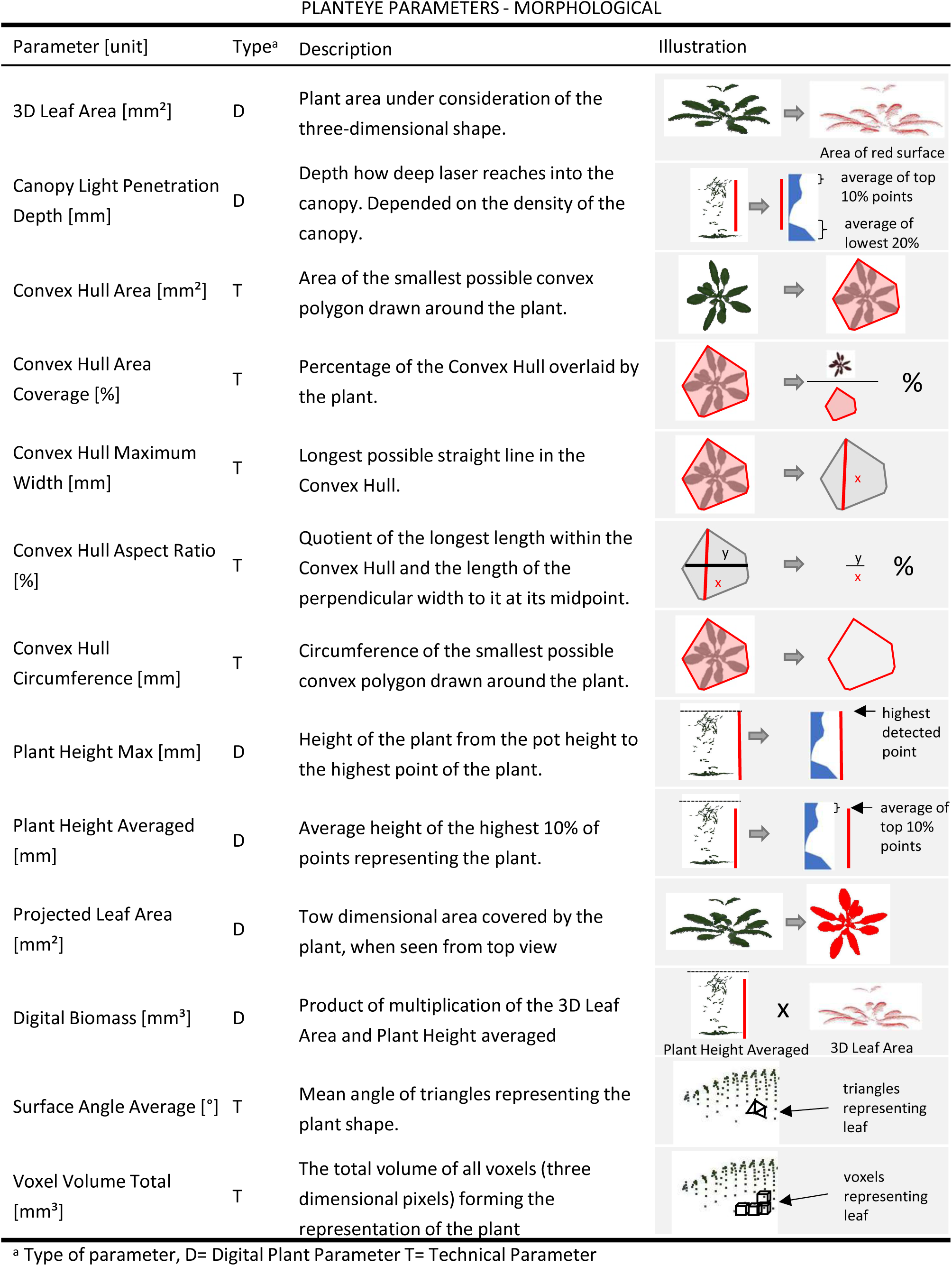
Morphological Machine-Derived Parameters Measured with the PlantEye. Thirteen different morphological parameters were measured at six time points across the life cycle. Unless indicated differently, the information on the parameters were obtained from the Phenospex-website and the manual provided by the company. The morphological parameters rely on the laser reflectance of an object. Digital Plant Parameters (here Type D) and Technical Parameters (here Type T) can be distinguished. Phena version Phena v2.0 and HortControl version 3.8.5 were used.

**Supplemental Figure 5:**
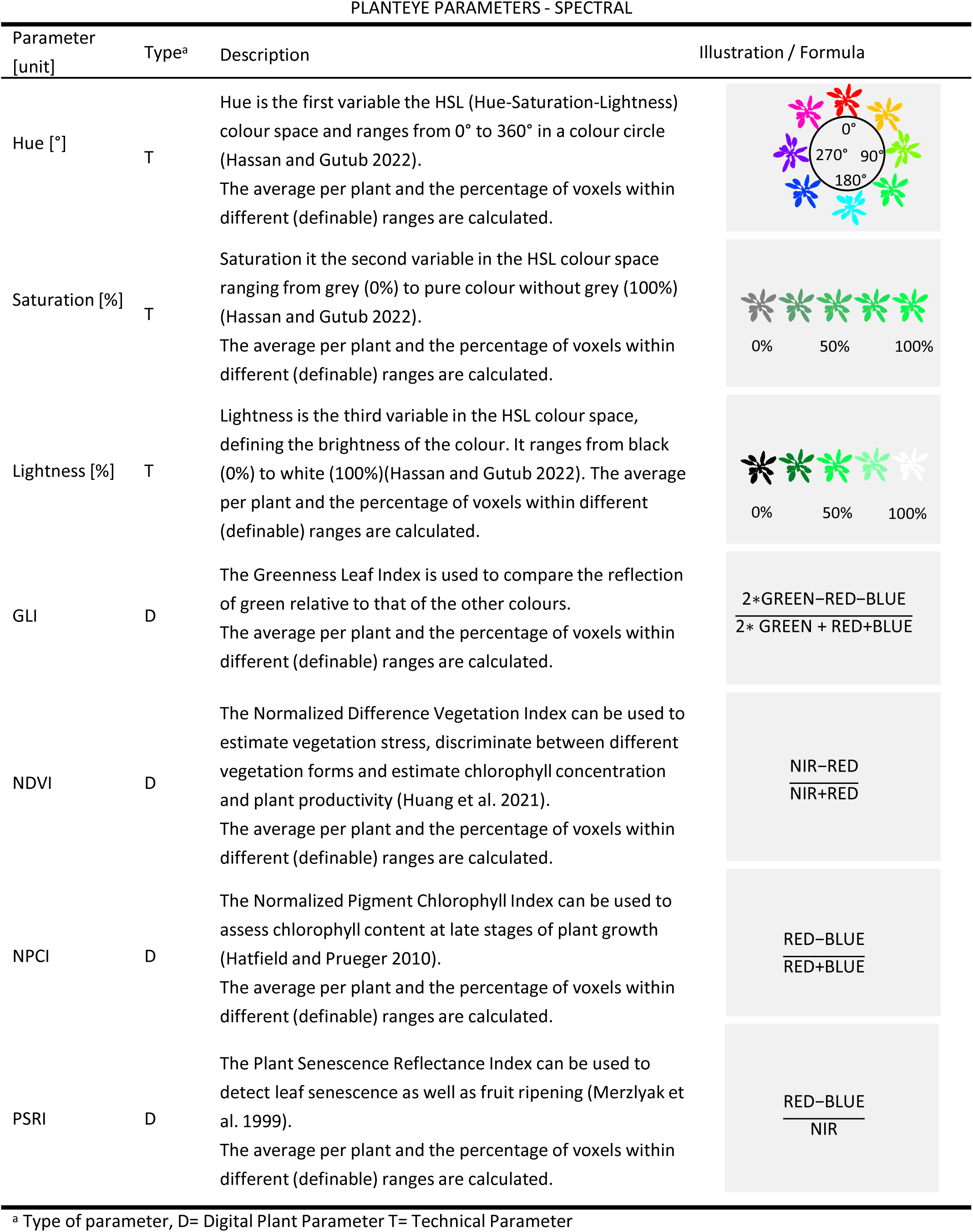
Spectral Machine-Derived Parameters Measured with the PlantEye. Unless indicated differently, the information on the parameters were obtained from the Phenospex-website and the manual provided by the company. Seven different spectral parameters can be determined. The colors in capital letters indicate the reflections of RED (624-634 nm), GREEN (530-540 nm), BLUE (465-485 nm) and NEAR-INFRARED (720-750 nm). For each parameter the average over the whole plant and the percentage of voxels within ranges (bins) is calculated. Digital Plant Parameters (here Type D) and Technical Parameters (here Type T) can be distinguished. Phena version Phena v2.0 and HortControl version 3.8.5 were used.

**Supplemental Figure 6:**
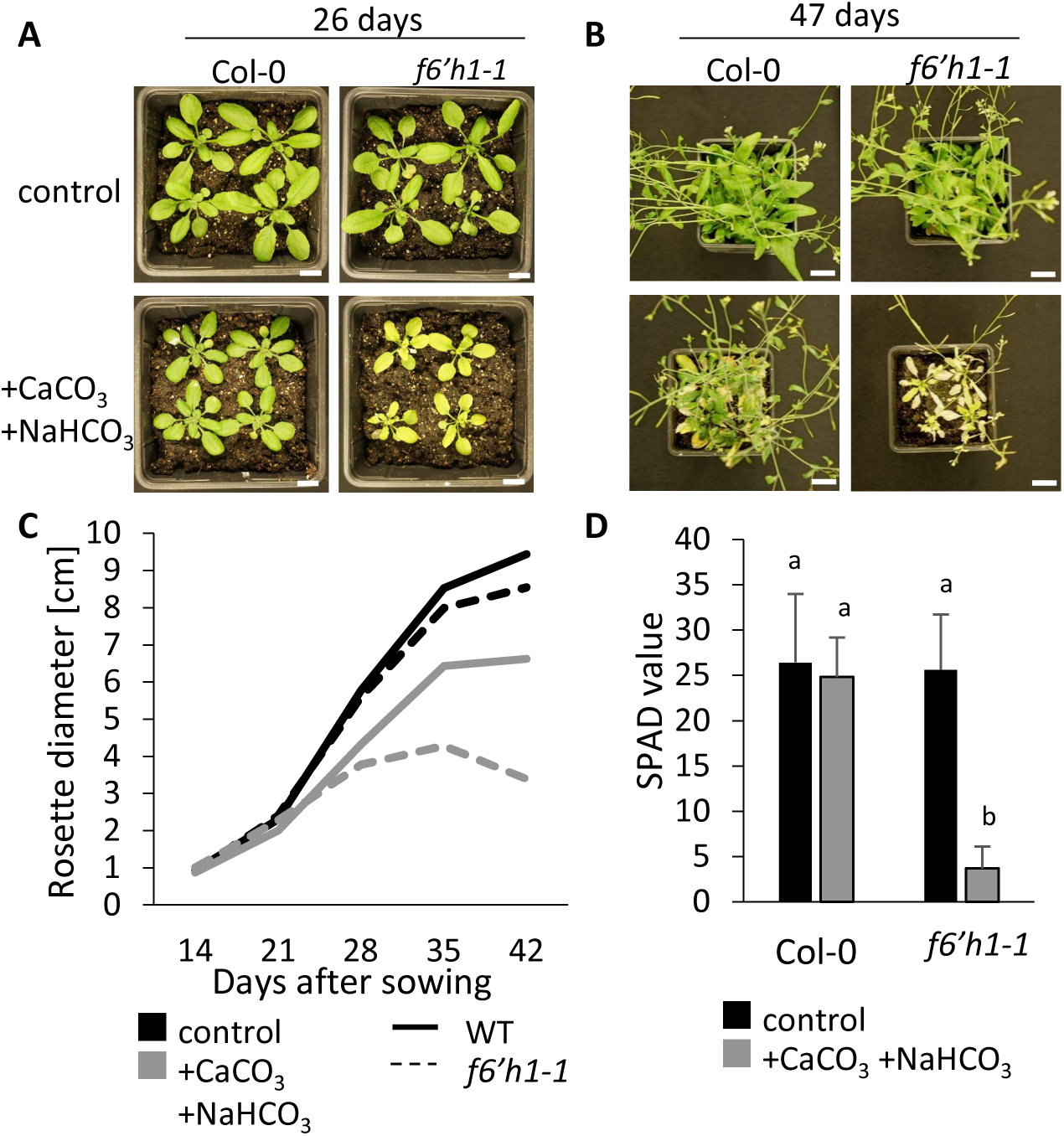
Preliminary Experiment Relying on Manual Phenotyping to Distinguish Wild Type and f6’h1-1 Mutants. In a preliminary experiment, we confirmed that indeed growth in an alkaline calcareous soil (ACS) condition causes reduced size of WT and *f6’h1-1* with more intense visible leaf chlorosis in *f6’h1-1*, confirmed by manual measurements of rosette diameter and SPAD values. **A, B:** Photos of Col-0 and *f6’h1-1* plants in control soil and soil with CaCO_3_ and NaHCO_3_ added 26 and 47 days after sowing. Scale bar=1 cm. C: Development of the rosette diameter in that condition (N= 16). **D:** SPAD values of Col-0 and *f6’h1-1* plants in control soil and soil with CaCO_3_ and NaHCO_3_ added 47/48 days after sowing. Labels indicate significantly different groups (p=0.05, Two-way ANOVA and Tukey Test in R, N=16 plants).

**Supplemental Figure 7:**
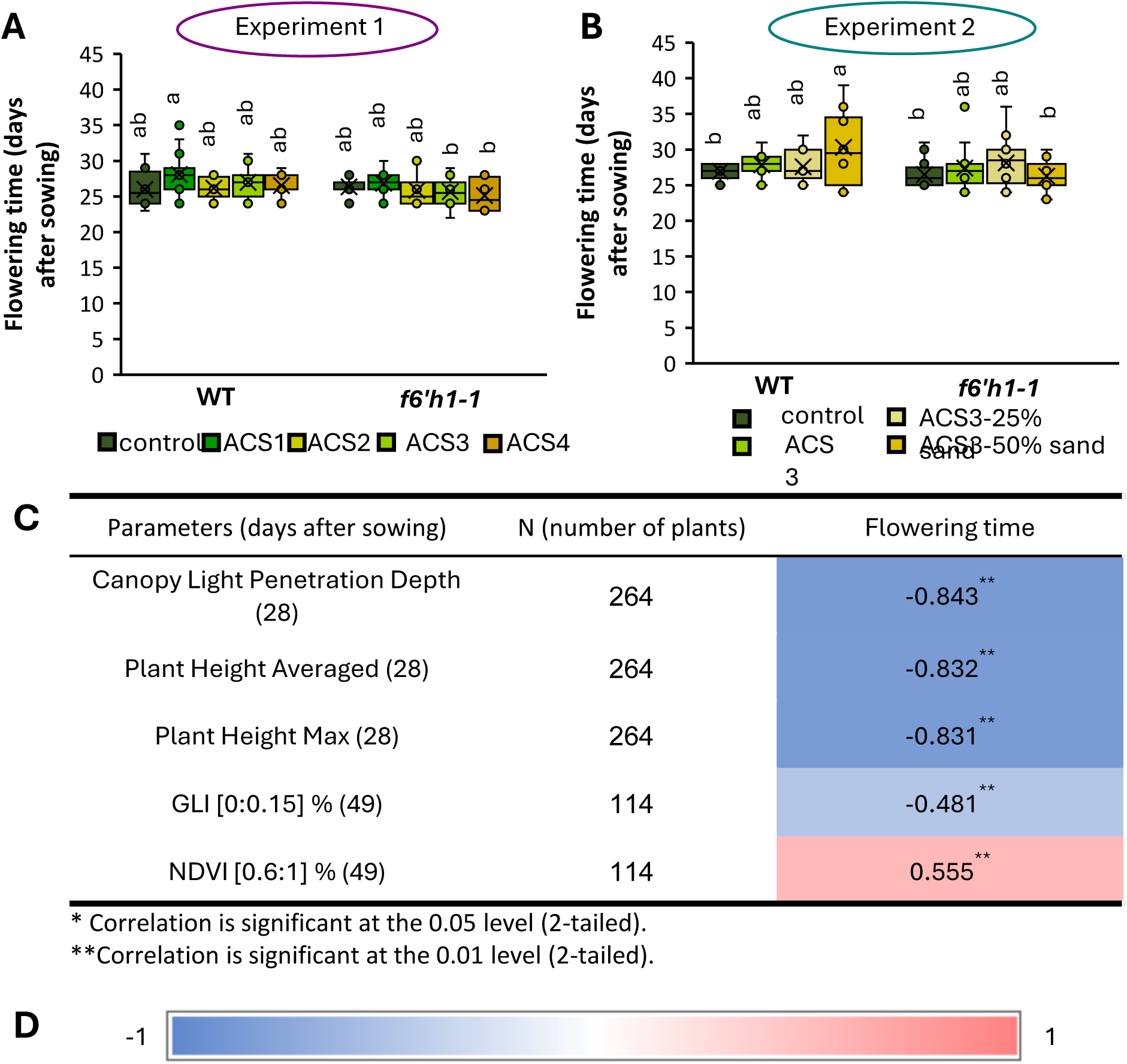
Flowering Time of Wild Type and *f6’h1-1* Mutant in Different Alkaline Calcareous (ACS) Conditions and Correlation with Machine-Derived Parameters (PlantEye). **A**: Flowering time of wildtype (WT) and *f6’h1-1* in control condition and conditions ACS1-4 (Experiment 1). **B:** Flowering time of WT and *f6’h1-1* in control condition and conditions ACS3, ACS3-25% sand and ACS3-50% sand (Experiment 2). Labels indicate statistical groups (two-way ANOVA and Tukey test in R, p=0.05, N=8-16). **C**: Machine-derived parameters correlating strongest (Spearman Rho correlation coefficient) with the flowering time. **D:** Color scale of heat map (-1 blue, 0 white, +1 red).

**Supplemental Figure 8:**
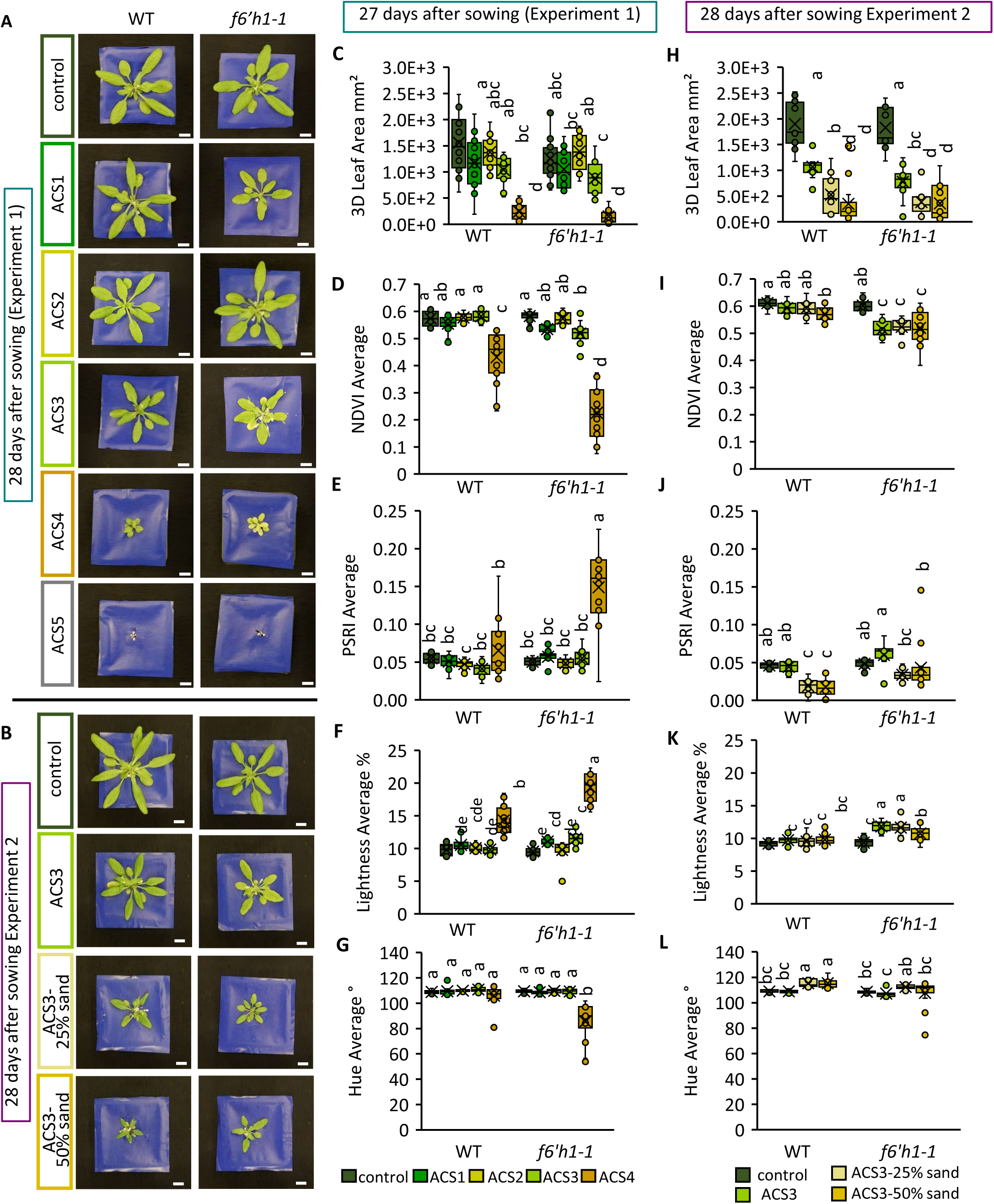
Machine-Derived (PlantEye) Phenotypical Analysis of Wild Type (WT) and *f6’h1-1* in Seven Alkaline Calcareous Soil (ACS) Conditions 27/28 days after sowing (DAS). *A. thaliana* wildtype (WT) and the coumarin deficient mutant *f6’h1-1 were* grown in seven alkaline calcareous soil (ACS) conditions with varying pH values in two experiments to determine an intermediate condition differentiating the lines. **A:** Plants in ACS1-5 (experiment 1) 28 days after sowing. Scale bar =1 cm. **B:** Plants in ACS3, ACS3-25% sand and ACS3 50%-sand (experiment 2) 28 days after sowing. Scale bar = 1 cm. **C-G:** 3D Leaf Area, Normalized Difference Vegetation Index (NDVI) Average, Plant Senescence Reflectance Index (PSRI) Average, Lightness Average and Hue Average in experiment 1 27 days after sowing determined with the PlantEye. **H-L:** 3D Leaf Area, Normalized Difference Vegetation Index (NDVI) Average, Plant Senescence Reflectance Index (PSRI) Average, Lightness Average and Hue Average in experiment 2 28 days after sowing determined with the PlantEye. Letters indicate statistical significance. N=9-16. Two-way ANOVA and Tukey-Test were performed in R. *p*=0.05. Graphs were created in Excel.

**Supplemental Table 1:**
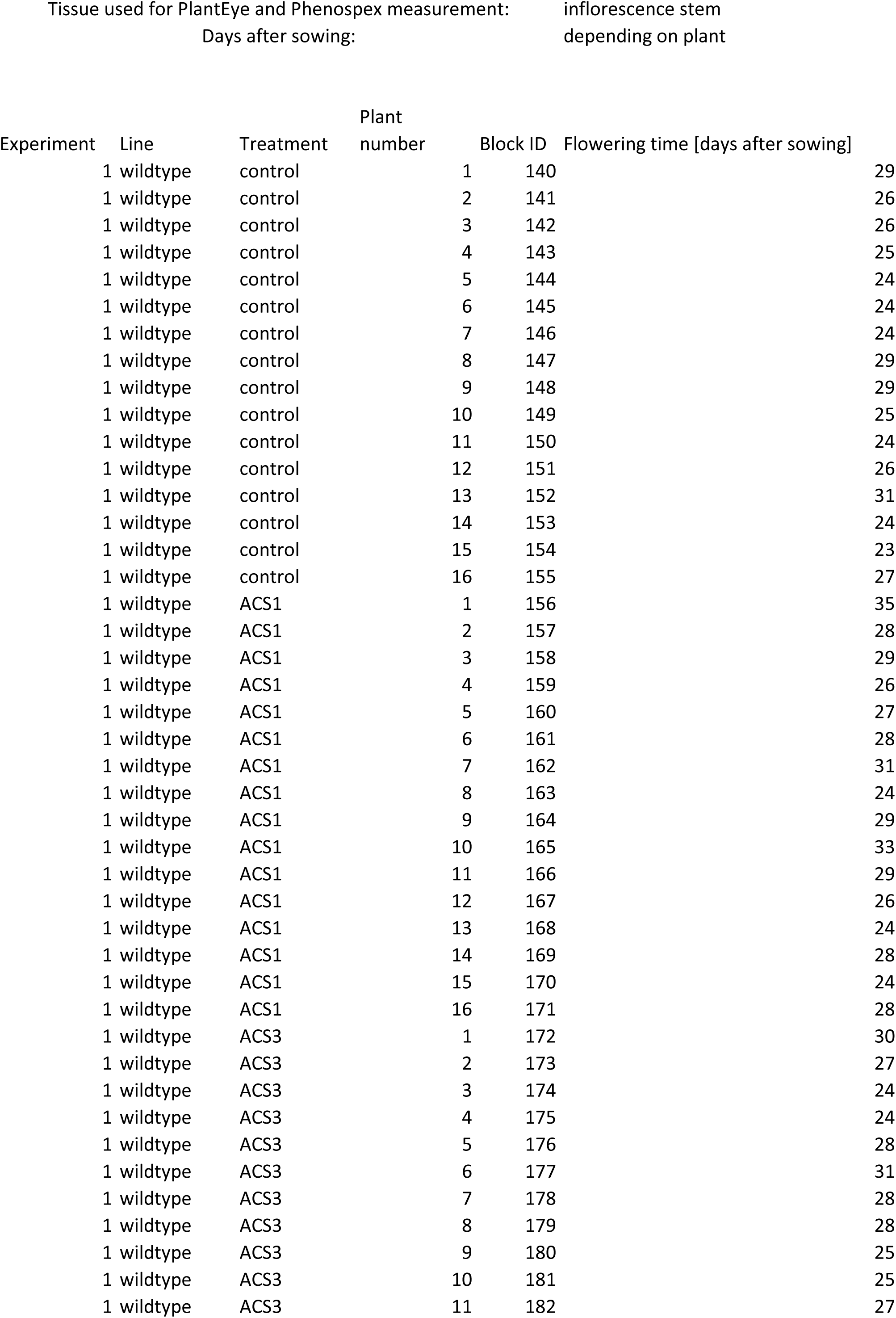
All Phenotypic Data Collected Manually and Machine-aided (PlantEye) for Correlation Analysis. Data of twelve suitable manual and 20 machine-derived PlantEye parameters for A*. thaliana* wildtype (WT) and its coumarin-deficient mutant *f6’h1-1* under control and up to seven alkaline calcareous soil (ACS) conditions, representing a scale of differing pH values from pH 6.2 (control) up to 8.3 (severe ACS) were recorded during two experiments. Manual parameters and PlantEye were determined at the indicated time points in days after sowing (DAS). Both whole plants or rosettes only were used for the determination of the parameters as indicated in the table. For details on the parameters see the material and methods section. For the PlantEye data, averages of repeated measurements per time point are shown. Block ID of plants are unique identifiers within the experiments. In case of empty fields, data were not available at for the parameters at the time point for the plant.

**Supplemental Table 2:**
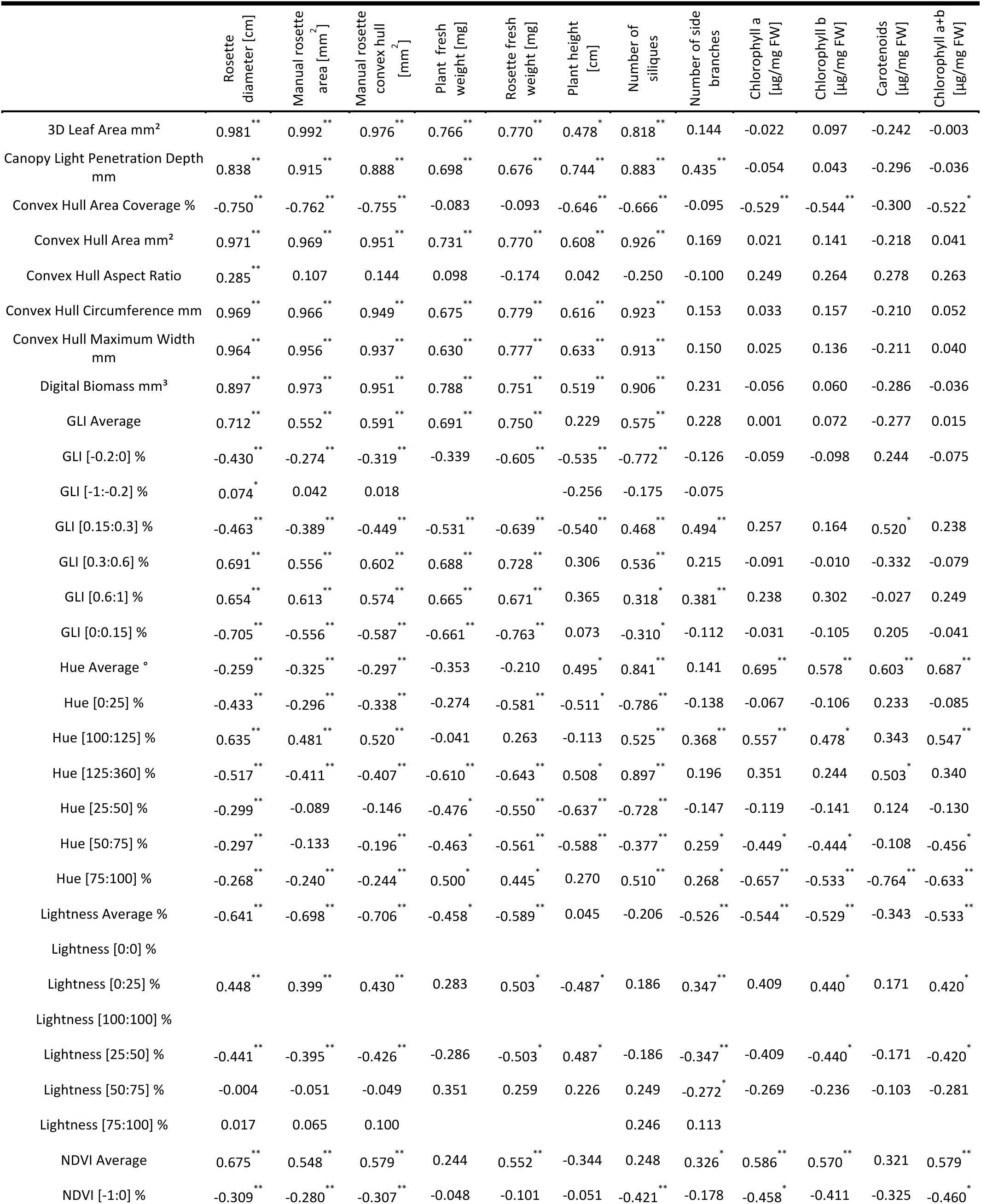
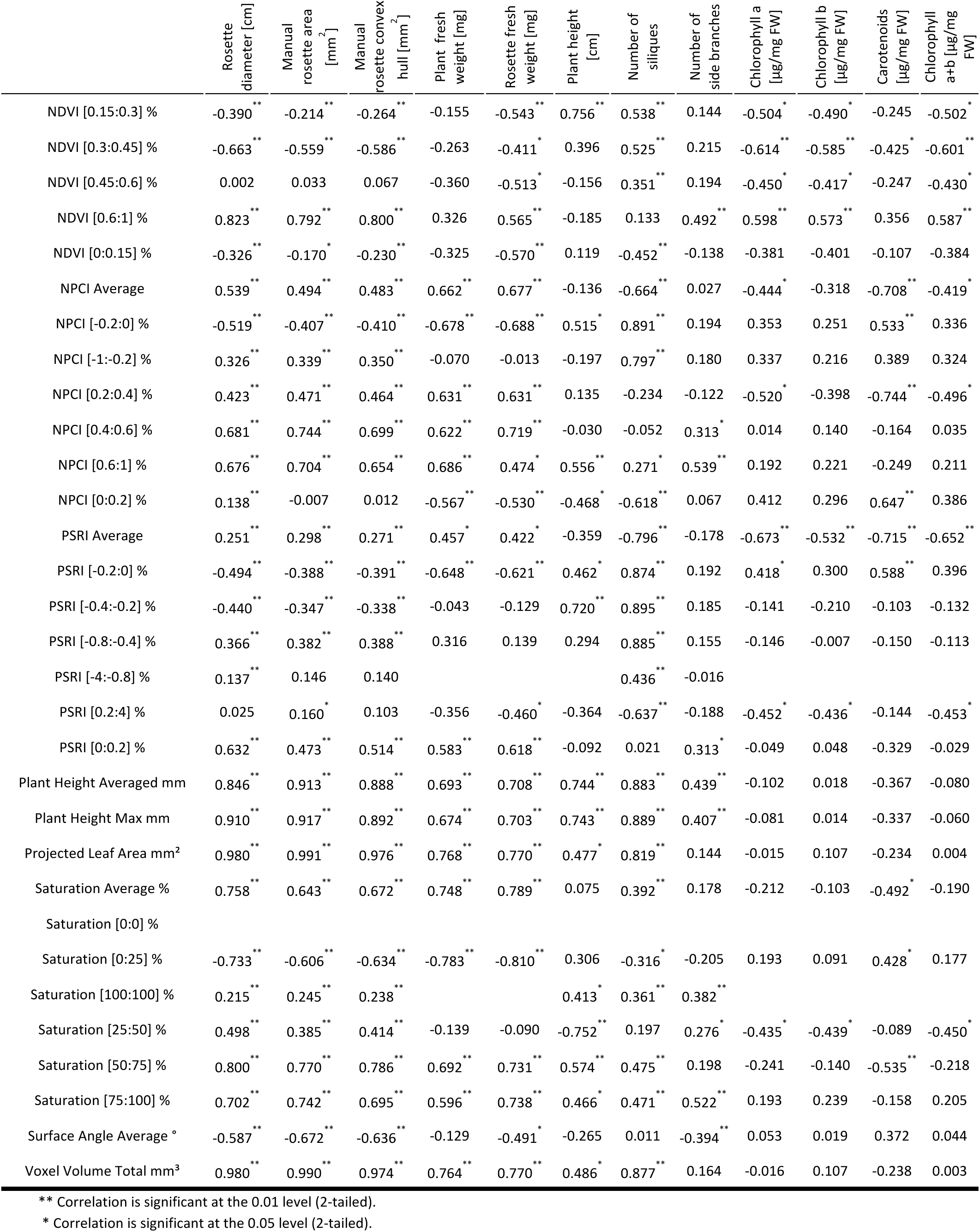
Results of Correlation Analysis of All Manually Determined and All Machine-Derived PlantEye Parameters. Data of twelve meaningful manual and 20 machine-derived PlantEye parameters for *A. thaliana* wildtype (WT) and its coumarin-deficient mutant *f6’h1-1* under control and up to seven alkaline calcareous soil (ACS) conditions, representing a scale of differing pH values from pH 6.2 (control) up to 8.3 (severe ACS) were recorded during two experiments and subjected to correlation analysis. Plant weight, rosette fresh weight and pigment contents were determined 38 days after sowing. Plant height was determined 44 days after sowing. Rosette diameter, manual rosette area and manual rosette convex hull were measured weekly during six weeks. N depended on parameters, if several time points were measured, plants were measured repeatedly. N (plant weight, rosette fresh weight, plant height) =24, N (Rosette diameter) =799, N (manual rosette area, manual convex hull area) =179, N (Number of siliques and side branches) = 61, N (pigment contents) = 23. Only rosettes were used for chlorophyll content measurement in acetone. For details on parameters see materials and methods section. Spearman Rho Correlation was done in SPSS. Significant correlations are marked (* <0.05, ** <0.01).

|  | Rosette diameter [cm] | Manual rosette area [mm<sup>2</sup>] | Manual rosette convex hull [mm<sup>2</sup>] | Plant fresh weight [mg] | Rosette fresh weight [mg] | Plant height [cm] | Number of siliques | Number of side branches | Chlorophyll a [µg/mg FW] | Chlorophyll b [µg/mg FW] | Carotenoids [µg/mg FW] | Chlorophyll a+b [µg/mg FW] |
| --- | --- | --- | --- | --- | --- | --- | --- | --- | --- | --- | --- | --- |
| 3D Leaf Area mm<sup>2</sup> | 0.981** | 0.992** | 0.976** | 0.766** | 0.770** | 0.478* | 0.818** | 0.144 | -0.022 | 0.097 | -0.242 | -0.003 |
| Canopy Light Penetration Depth mm | 0.838** | 0.915** | 0.888** | 0.698** | 0.676** | 0.744** | 0.883** | 0.435** | -0.054 | 0.043 | -0.296 | -0.036 |
| Convex Hull Area Coverage % | -0.750** | -0.762** | -0.755** | -0.083 | -0.093 | -0.646** | -0.666** | -0.095 | -0.529** | -0.544** | -0.300 | -0.522* |
| Convex Hull Area mm<sup>2</sup> | 0.971** | 0.969** | 0.951** | 0.731** | 0.770** | 0.608** | 0.926** | 0.169 | 0.021 | 0.141 | -0.218 | 0.041 |
| Convex Hull Aspect Ratio | 0.285** | 0.107 | 0.144 | 0.098 | -0.174 | 0.042 | -0.250 | -0.100 | 0.249 | 0.264 | 0.278 | 0.263 |
| Convex Hull Circumference mm | 0.969** | 0.966** | 0.949** | 0.675** | 0.779** | 0.616** | 0.923** | 0.153 | 0.033 | 0.157 | -0.210 | 0.052 |
| Convex Hull Maximum Width mm | 0.964** | 0.956** | 0.937** | 0.630** | 0.777** | 0.633** | 0.913** | 0.150 | 0.025 | 0.136 | -0.211 | 0.040 |
| Digital Biomass mm<sup>3</sup> | 0.897** | 0.973** | 0.951** | 0.788** | 0.751** | 0.519** | 0.906** | 0.231 | -0.056 | 0.060 | -0.286 | -0.036 |
| GLI Average | 0.712** | 0.552** | 0.591** | 0.691** | 0.750** | 0.229 | 0.575** | 0.228 | 0.001 | 0.072 | -0.277 | 0.015 |
| GLI [-0.2:0] % | -0.430** | -0.274** | -0.319** | -0.339 | -0.605** | -0.535** | -0.772** | -0.126 | -0.059 | -0.098 | 0.244 | -0.075 |
| GLI [-1:-0.2] % | 0.074* | 0.042 | 0.018 |  |  | -0.256 | -0.175 | -0.075 |  |  |  |  |
| GLI [0.15:0.3] % | -0.463** | -0.389** | -0.449** | -0.531** | -0.639** | -0.540** | 0.468** | 0.494** | 0.257 | 0.164 | 0.520* | 0.238 |
| GLI [0.3:0.6] % | 0.691** | 0.556** | 0.602** | 0.688** | 0.728** | 0.306 | 0.536** | 0.215 | -0.091 | -0.010 | -0.332 | -0.079 |
| GLI [0.6:1] % | 0.654** | 0.613** | 0.574** | 0.665** | 0.671** | 0.365 | 0.318* | 0.381** | 0.238 | 0.302 | -0.027 | 0.249 |
| GLI [0:0.15] % | -0.705** | -0.556** | -0.587** | -0.661** | -0.763** | 0.073 | -0.310* | -0.112 | -0.031 | -0.105 | 0.205 | -0.041 |
| Hue Average ° | -0.259** | -0.325** | -0.297** | -0.353 | -0.210 | 0.495* | 0.841** | 0.141 | 0.695** | 0.578** | 0.603** | 0.687** |
| Hue [0:25] % | -0.433** | -0.296** | -0.338** | -0.274 | -0.581** | -0.511* | -0.786** | -0.138 | -0.067 | -0.106 | 0.233 | -0.085 |
| Hue [100:125] % | 0.635** | 0.481** | 0.520** | -0.041 | 0.263 | -0.113 | 0.525** | 0.368** | 0.557** | 0.478* | 0.343 | 0.547** |
| Hue [125:360] % | -0.517** | -0.411** | -0.407** | -0.610** | -0.643** | 0.508* | 0.897** | 0.196 | 0.351 | 0.244 | 0.503* | 0.340 |
| Hue [25:50] % | -0.299** | -0.089 | -0.146 | -0.476* | -0.550** | -0.637** | -0.728** | -0.147 | -0.119 | -0.141 | 0.124 | -0.130 |
| Hue [50:75] % | -0.297** | -0.133 | -0.196** | -0.463* | -0.561** | -0.588** | -0.377** | 0.259* | -0.449* | -0.444* | -0.108 | -0.456* |
| Hue [75:100] % | -0.268** | -0.240** | -0.244** | 0.500* | 0.445* | 0.270 | 0.510** | 0.268* | -0.657** | -0.533** | -0.764** | -0.633** |
| Lightness Average % | -0.641** | -0.698** | -0.706** | -0.458* | -0.589** | 0.045 | -0.206 | -0.526** | -0.544** | -0.529** | -0.343 | -0.533** |
| Lightness [0:0] % |  |  |  |  |  |  |  |  |  |  |  |  |
| Lightness [0:25] % | 0.448** | 0.399** | 0.430** | 0.283 | 0.503* | -0.487* | 0.186 | 0.347** | 0.409 | 0.440* | 0.171 | 0.420* |
| Lightness [100:100] % |  |  |  |  |  |  |  |  |  |  |  |  |
| Lightness [25:50] % | -0.441** | -0.395** | -0.426** | -0.286 | -0.503* | 0.487* | -0.186 | -0.347** | -0.409 | -0.440* | -0.171 | -0.420* |
| Lightness [50:75] % | -0.004 | -0.051 | -0.049 | 0.351 | 0.259 | 0.226 | 0.249 | -0.272* | -0.269 | -0.236 | -0.103 | -0.281 |
| Lightness [75:100] % | 0.017 | 0.065 | 0.100 |  |  |  | 0.246 | 0.113 |  |  |  |  |
| NDVI Average | 0.675** | 0.548** | 0.579** | 0.244 | 0.552** | -0.344 | 0.248 | 0.326* | 0.586** | 0.570** | 0.321 | 0.579** |
| NDVI [-1:0] % | -0.309** | -0.280** | -0.307** | -0.048 | -0.101 | -0.051 | -0.421** | -0.178 | -0.458* | -0.411 | -0.325 | -0.460* |
| NDVI [0.15:0.3] % | -0.390** | -0.214** | -0.264** | -0.155 | -0.543** | 0.756** | 0.538** | 0.144 | -0.504* | -0.490* | -0.245 | -0.502* |
| NDVI [0.3:0.45] % | -0.663** | -0.559** | -0.586** | -0.263 | -0.411* | 0.396 | 0.525** | 0.215 | -0.614** | -0.585** | -0.425* | -0.601** |
| NDVI [0.45:0.6] % | 0.002 | 0.033 | 0.067 | -0.360 | -0.513* | -0.156 | 0.351** | 0.194 | -0.450* | -0.417* | -0.247 | -0.430* |
| NDVI [0.6:1] % | 0.823** | 0.792** | 0.800** | 0.326 | 0.565** | -0.185 | 0.133 | 0.492** | 0.598** | 0.573** | 0.356 | 0.587** |
| NDVI [0:0.15] % | -0.326** | -0.170* | -0.230** | -0.325 | -0.570** | 0.119 | -0.452** | -0.138 | -0.381 | -0.401 | -0.107 | -0.384 |
| NPCI Average | 0.539** | 0.494** | 0.483** | 0.662** | 0.677** | -0.136 | -0.664** | 0.027 | -0.444* | -0.318 | -0.708** | -0.419* |
| NPCI [-0.2:0] % | -0.519** | -0.407** | -0.410** | -0.678** | -0.688** | 0.515* | 0.891** | 0.194 | 0.353 | 0.251 | 0.533** | 0.336 |
| NPCI [-1:-0.2] % | 0.326** | 0.339** | 0.350** | -0.070 | -0.013 | -0.197 | 0.797** | 0.180 | 0.337 | 0.216 | 0.389 | 0.324 |
| NPCI [0.2:0.4] % | 0.423** | 0.471** | 0.464** | 0.631** | 0.631** | 0.135 | -0.234 | -0.122 | -0.520* | -0.398 | -0.744** | -0.496* |
| NPCI [0.4:0.6] % | 0.681** | 0.744** | 0.699** | 0.622** | 0.719** | -0.030 | -0.052 | 0.313* | 0.014 | 0.140 | -0.164 | 0.035 |
| NPCI [0.6:1] % | 0.676** | 0.704** | 0.654** | 0.686** | 0.474* | 0.556** | 0.271* | 0.539** | 0.192 | 0.221 | -0.249 | 0.211 |
| NPCI [0:0.2] % | 0.138** | -0.007 | 0.012 | -0.567** | -0.530** | -0.468* | -0.618** | 0.067 | 0.412 | 0.296 | 0.647** | 0.386 |
| PSRI Average | 0.251** | 0.298** | 0.271** | 0.457* | 0.422* | -0.359 | -0.796** | -0.178 | -0.673** | -0.532** | -0.715** | -0.652** |
| PSRI [-0.2:0] % | -0.494** | -0.388** | -0.391** | -0.648** | -0.621** | 0.462* | 0.874** | 0.192 | 0.418* | 0.300 | 0.588** | 0.396 |
| PSRI [-0.4:-0.2] % | -0.440** | -0.347** | -0.338** | -0.043 | -0.129 | 0.720** | 0.895** | 0.185 | -0.141 | -0.210 | -0.103 | -0.132 |
| PSRI [-0.8:-0.4] % | 0.366** | 0.382** | 0.388** | 0.316 | 0.139 | 0.294 | 0.885** | 0.155 | -0.146 | -0.007 | -0.150 | -0.113 |
| PSRI [-4:-0.8] % | 0.137** | 0.146 | 0.140 |  |  |  | 0.436** | -0.016 |  |  |  |  |
| PSRI [0.2:4] % | 0.025 | 0.160* | 0.103 | -0.356 | -0.460* | -0.364 | -0.637** | -0.188 | -0.452* | -0.436* | -0.144 | -0.453* |
| PSRI [0:0.2] % | 0.632** | 0.473** | 0.514** | 0.583** | 0.618** | -0.092 | 0.021 | 0.313* | -0.049 | 0.048 | -0.329 | -0.029 |
| Plant Height Averaged mm | 0.846** | 0.913** | 0.888** | 0.693** | 0.708** | 0.744** | 0.883** | 0.439** | -0.102 | 0.018 | -0.367 | -0.080 |
| Plant Height Max mm | 0.910** | 0.917** | 0.892** | 0.674** | 0.703** | 0.743** | 0.889** | 0.407** | -0.081 | 0.014 | -0.337 | -0.060 |
| Projected Leaf Area mm<sup>2</sup> | 0.980** | 0.991** | 0.976** | 0.768** | 0.770** | 0.477* | 0.819** | 0.144 | -0.015 | 0.107 | -0.234 | 0.004 |
| Saturation Average % | 0.758** | 0.643** | 0.672** | 0.748** | 0.789** | 0.075 | 0.392** | 0.178 | -0.212 | -0.103 | -0.492* | -0.190 |
| Saturation [0:0] % |  |  |  |  |  |  |  |  |  |  |  |  |
| Saturation [0:25] % | -0.733** | -0.606** | -0.634** | -0.783** | -0.810** | 0.306 | -0.316* | -0.205 | 0.193 | 0.091 | 0.428* | 0.177 |
| Saturation [100:100] % | 0.215** | 0.245** | 0.238** |  |  | 0.413* | 0.361** | 0.382** |  |  |  |  |
| Saturation [25:50] % | 0.498** | 0.385** | 0.414** | -0.139 | -0.090 | -0.752** | 0.197 | 0.276* | -0.435* | -0.439* | -0.089 | -0.450* |
| Saturation [50:75] % | 0.800** | 0.770** | 0.786** | 0.692** | 0.731** | 0.574** | 0.475** | 0.198 | -0.241 | -0.140 | -0.535** | -0.218 |
| Saturation [75:100] % | 0.702** | 0.742** | 0.695** | 0.596** | 0.738** | 0.466* | 0.471** | 0.522** | 0.193 | 0.239 | -0.158 | 0.205 |
| Surface Angle Average ° | -0.587** | -0.672** | -0.636** | -0.129 | -0.491* | -0.265 | 0.011 | -0.394** | 0.053 | 0.019 | 0.372 | 0.044 |
| Voxel Volume Total mm<sup>3</sup> | 0.980** | 0.990** | 0.974** | 0.764** | 0.770** | 0.486* | 0.877** | 0.164 | -0.016 | 0.107 | -0.238 | 0.003 |

\*\* Correlation is significant at the 0.01 level (2-tailed).
\* Correlation is significant at the 0.05 level (2-tailed).

**Supplemental Table 3:**
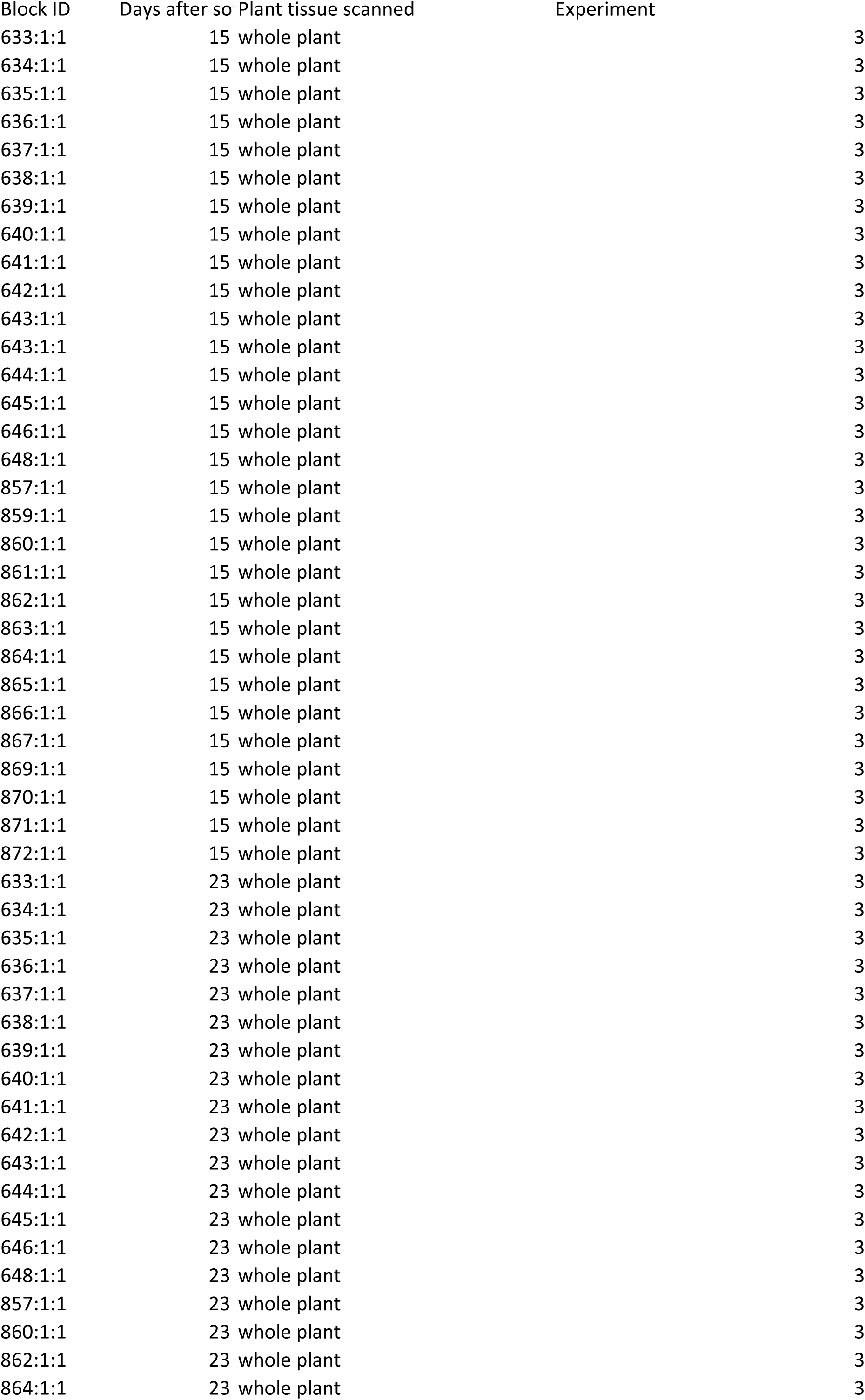
All Machine-Derived Data for Phenotypic Analysis Using Known Regulatory Iron Homeostasis Mutants. The ACS3 condition and the selected phenotyping parameters were applied to validate the procedure using iron (Fe) homeostasis mutants with the additional aim of identifying potentially novel phenotypes appearing during the life cycle. The iron homeostasis mutants *fit-3, pye-1* and *btsl1 btsl2* were phenotyped in ACS3 weekly. Eight plants per line were planted. The data were obtained at the time points indicated in the column days after sowing. The Block ID served as a unique identifier for each plant. Note that some mutant plants did not survive.

